# Integrative Transcriptomic and Machine Learning Analysis of ecDNA-Associated Features for Studying Chemotherapy Resistance in TNBC

**DOI:** 10.64898/2026.04.02.716106

**Authors:** Md. Iftehimul, Dipongkor Saha

**Author notes:** Correspondence to: Dipongkor Saha, DVM, PhD, Department of Biology, College of Science and Technology, North Carolina Agricultural and Technical State University, 1601 East Market Street (Hines Hall 300B), Greensboro, NC 27411, USA. Authors’ email addresses: Md. Iftehimul.

## Abstract

Extrachromosomal DNA (ecDNA) has emerged as a critical mediator of oncogene amplification and transcriptional dynamics in aggressive cancers, yet its contribution to chemotherapy resistance *in vivo* remains incompletely understood. This study investigates the contribution of ecDNA-associated molecular features to predictive chemotherapy resistance in TNBC. We analyzed RNA-seq data from 4T1 TNBC cells and 4T1 bulk tumors at different growth stages (1-, 3-, and 6-week) to identify differentially expressed ecDNA alterations. We then utilized molecular docking tools to predict ecDNA protein-drug interactions and employed machine learning (ML) models to predict ecDNA-associated therapeutic resistance. Our results revealed changes in global gene expression, including expression of ecDNA-associated genes, that continued over time, with significant molecular remodeling observed at six weeks. Additionally, we found gradual accumulation of mutations in ecDNA genes, which may have contributed to reduced drug binding affinity, indicating potential resistance. ML models generated stable, high-confidence classifications of resistant phenotypes, consistently identifying ecDNA burden and prevalence as dominant predictive features of drug resistance. Drug specific predictions further highlighted elevated resistance probabilities for paclitaxel and doxorubicin, whereas hydroxyurea, which depletes ecDNA, showed reduced resistance probabilities, indicating potential roles of ecDNA in chemoresistance. This study provides new insights into temporal remodeling of ecDNA within TNBC tumors over time and their potential association with drug resistance.

## 1. Introduction

Extrachromosomal DNAs (ecDNAs) are circular, non-chromosomal DNA elements (1-3 Mb)^1^ that are frequently found in tumor cells.^2,3^ EcDNA has been increasingly linked to oncogene amplification, elevated transcriptional output, and therapeutic resistance in cancer.^4,5^ Triple-negative breast cancer (TNBC), an aggressive type of breast cancer,^6^ provides a useful model system for studying ecDNA due to its reported high prevalence in breast neoplasms.^7^ While direct structural characterization of ecDNA requires specialized genomic and experimental approaches, genes previously reported to be ecDNA-associated in cancer contexts may display distinctive and dynamic transcriptional behavior during tumor progression.^8–15^ Characterizing such temporal transcriptional remodeling across tumor stages may offer insight into potential ecDNA-related mechanisms underlying chemotherapy resistance, an area that remains poorly understood.

Chemotherapeutic agents such as doxorubicin (DOX) and paclitaxel (PAC) are commonly used to treat TNBC.^16,17^ DOX, an anthracycline, intercalates DNA, inhibits topoisomerase II, and alters the cellular redox environment via mitochondrial redox cycling,^16,18^ whereas PAC, a taxane, stabilizes microtubules and disrupts mitotic progression.^17^ Despite their clinical utility, resistance to these agents is frequently observed in TNBC. Since ecDNA has been implicated as a potential contributor to resistance-associated phenotypes in several cancer contexts,^2,10,12,14,19^ we wanted to investigate whether ecDNA-associated molecular features correlate with predicted resistance behavior.

In this study, we aimed to characterize global gene expression dynamics, focusing specifically on temporal molecular remodeling of ecDNA-associated genes, across distinct stages of *in vivo* tumor growth in the 4T1 TNBC model. We further sought to examine whether transcript-derived sequence variation inferred from RNA-seq data in selected ecDNA-related genes varies across tumor progression. However, the high dimensionality and complexity of transcriptomic datasets present challenges in predicting ecDNA-associated genes linked to chemoresistance.^20^ Machine learning (ML) offers a robust solution, enabling the integration of ecDNA features with gene expression data to detect subtle patterns, classify resistant versus sensitive phenotypes, and predict drug response.^21^ To address this, we applied supervised learning algorithms, including Random Forest,^22^ XGBoost,^23^ and Logistic Regression,^24,25^ to evaluate whether ecDNA-associated molecular features are linked to chemoresistance. Consistent with these objectives, we observed dynamic global transcriptional changes across tumor stages, including temporal remodeling of ecDNA-associated genes across time points with increased transcript-derived sequence variation in selected ecDNA genes. In addition, ML models predicted ecDNA-associated genes as potential drivers of chemotherapy resistance.

## 2. Methodology

### 2.1. Datasets acquisition and statistical analysis of differentially expressed genes (DEGs)

The 4T1 murine tumor cell line is widely used as a TNBC model to study tumor progression and treatment response.^26^ To determine temporal molecular remodeling of ecDNA, we extracted RNA-seq datasets (originating from 4T1 cells as a baseline and 1-week-, 3-week-, and 6-week-old 4T1 tumors derived from the same batch of 4T1 cells) from the NCBI database (***Table S1***). The quality of raw sequencing reads was assessed using FastQC (v.0.12.1).^27^ Low-quality bases and adapter sequences were removed using Trimmomatic (v0.4)^28^ and aligned the reads to the GRCm39 (Ensembl release 115) genome using HISAT2.^29^ *De novo* assembly was performed using Trinity (v2.15).^30^ We quantified gene-level read counts using FeatureCounts,^31^ normalized them across samples using DESeq2,^32^ and analyzed DEGs. Genes with adjusted p-value <0.05 and Log2FC ≥ 1 were considered to be potential DEGs. Additionally, we used Log2FC ≥ 1 and Log2FC ≤ −1 criteria to explore up- and down-regulated genes, respectively. DEGs were then imported into R Studio to generate the volcano plots and principal component analysis (PCA).^33^

### 2.2. Interaction of ecDNA-related proteins and anti-TNBC drugs

The three-dimensional structures of wild-type and mutated ecDNA-related oncoproteins were obtained from the Protein Data Bank. The protein structures were selected based on their crystallographic data, with metals, co-factors, water molecules, and side chains excluded. The active residues of these proteins were identified using the COACH-D algorithm. To enhance the specificity of the screening, chain A was selected and prepared by removing unwanted ions, heteroatoms, ligands, and water molecules. Additionally, hydrogen atoms were added to the proteins to ensure proper functioning during the screening process. All protein preparation steps were performed using the UCSF Chimera 1.17.3 platform.^34^

The molecular screening study was conducted using the PyRx 0.8 package,^35^ a computational molecular docking toolkit. A grid box was created within the active site of the core protein, with dimensions x = 28.33, y = 35.10, and z = 39.48. The grid center was positioned at coordinates x = 26.72, y= −10.14, and z = 1.45.^36^ The grid box position was kept constant for each selected core protein to maintain a consistent screening environment. AutoDock Vina was run via a batch script, which sequentially executed commands to operate the software.^37^ Following the docking process, multiple models (usually between 1 and 9) were generated for each receptor-ligand interaction. The model with the highest binding free energy and the lowest Root Mean Square Deviation (RMSD) score was identified as the optimal one.^38^ Hydrophobic and hydrogen bond interactions were further highlighted in the molecular interaction analysis using BIOVIA Discovery Studio.^39^

### 2.3. Machine Learning Based Prediction of Drug Resistance

#### 2.3.1. Data Preprocessing

The dataset comprising 40 experimental observations was compiled through systematic literature review. Data preprocessing was performed to ensure consistency and prevent modeling bias. All categorical text fields were standardized by removing encoding artifacts, trimming whitespace, and converting entries to lowercase. The primary outcome variable, drug resistance, was encoded as a binary variable (1 = resistant; 0 = sensitive) based on standardized textual mapping, and rows with undefined resistance status were excluded. Ordinal ecDNA-related descriptors, including ecDNA prevalence, were converted into numerical scores (high = 3, moderate = 2, low = 1) to preserve biological ranking while enabling ML compatibility and leakage-prone variables were excluded to ensure unbiased modeling. The dataset was then reshaped from wide to long format using a melt transformation, where each gene amplification event per sample became a separate observation. Through reshaping, gene amplification columns were consolidated into two variables: gene name and amplification status, enabling each gene amplification event per sample to be represented as a separate observation. Let *G* represent the number of amplification genes and *N* the number of samples; the total number of gene-level observations after reshaping was calculated as:

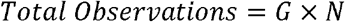

This transformation converted each gene amplification event into a separate observation for model training. As a result, the dataset expanded from 40 original samples to 1,160 observations, substantially increasing the number of training instances available for downstream ML-based prediction of drug resistance.

#### 2.3.2. Ideal feature finding

Feature selection was performed to identify the most relevant and informative variables for predicting ecDNA-related drug resistance using supervised ML models. This process reduces dimensionality, minimizes overfitting, and improves model interpretability by focusing on biologically meaningful features. In this study, we applied feature selection techniques including linear discriminant analysis (LDA) to identify the most informative features for ML-based models aiming to enhance the prediction of drug resistance.^40^

#### 2.3.3. Model Development

In this study, we applied a number of ML-based models to predict ecDNA-associated drug resistance and evaluated their performance using multiple parameters. These models are Logistic Regression (LR), Random Forest (RF), Extreme Gradient Boosting (XGBoost), and voting ensemble.

##### Logistic Regression (LR)

LR was employed as a baseline linear classifier,^24,25^ modeling the probability of a binary outcome *y* = ∈ {0,1} based on input features *x* = [*x*_1_, *x*_2_, … *x*_*p*_]. The relationship between features and the log-odds of resistance is given by:

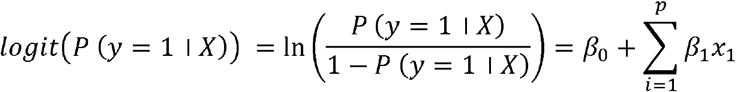

The predicted probability is obtained via sigmoid function:

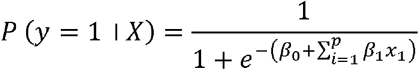

Here, *β*_0_ is the intercept and *β*_*i*_ is the feature coefficients. Features with higher | *β*_*i*_ | indicate stronger influence on resistance prediction.

##### Random Forest (RF)

RF is a non-parametric ensemble method that combines multiple decision trees to improve predictive performance and reduce variance.^22^ Each tree *T*_*j*_ is trained on a bootstrap sample of the data, and at each split, a random subset of features is considered. The predicted class is determined by majority voting across *M* trees:

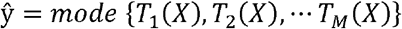

RF is robust to multicollinearity and capable of capturing nonlinear interactions between features, such as combinations of gene amplifications and drug properties that drive resistance. Feature importance scores were calculated based on the reduction in Gini impurity contributed by each variable across all trees, allowing identification of ecDNA-associated genes that strongly influence resistance.

#### Extreme Gradient Boosting (XGBoost)

XGBoost builds an additive model of *K* sequential decision trees *f*_*k*_ (*X*) to minimize a regularized objective function.^23^ The model predicts the outcome as:

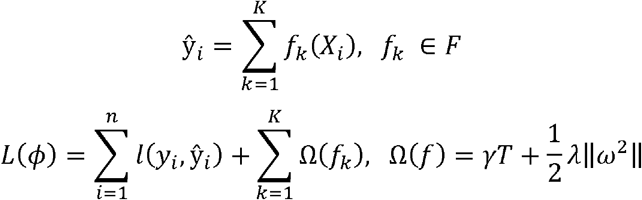

Here, *l* (*ŷ*_*i*_, *y*_*i*_) is the binary logistic loss, Ω (*f*_*k*_) is a regularization term to penalize model complexity, *T* is the number of leaves in a tree, Ω is the vector of leaf scores, and γ, λ are regularization parameters.

To further enhance predictive robustness and reduce model-specific bias, an ensemble learning strategy was implemented using a soft voting classifier.^41^ This approach integrates the predictive strengths of LR, RF, and XGBoost by aggregating their probabilistic outputs. The ensemble model was subsequently evaluated using accuracy, precision, recall, F1-score, log loss metrics, and receiver operating characteristic (ROC) with area under the curve (AUC) metrics. By combining complementary learning mechanisms, the ensemble framework provided a more balanced and reliable prediction of ecDNA-associated drug resistance. The final prediction is determined by averaging these probabilities and assigning the class with the highest aggregated probability:

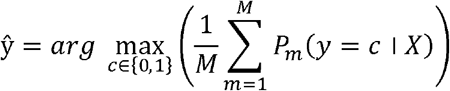

Here, *M* represents the number of base models (here, *M*=3) and *P*_*m*_ (*y* =*c*|*X*) denotes the predicted probability of class *c* from model *m*.

Finally, the quality and calibration of predicted probabilities were evaluated using learning curve analysis of log loss, which penalizes incorrect and overconfident predictions more heavily than simple accuracy.^42^ This matric is particularly important in drug resistance prediction, where probabilistic results may inform biological interpretation and therapeutic decision-making. For a binary classification problem, the log loss function is defined as:

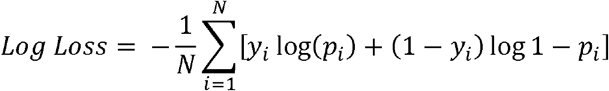

Here *N* is the number of samples, *y*_*i*_ represents the true class label and *p*_*i*_ denotes the predicted probability of resistance.

#### 2.3.4. Post processing and model validation

The developed ML models for predicting ecDNA-associated drug resistance were evaluated using 80:20 train-test split to ensure robust model training and realistic performance. Model predictions were analyzed using confusion matrix (CM) comprising True Positives (TP), False Positives (FP), and False Negatives (FN).^43^ Performance metrics were calculated as follows:

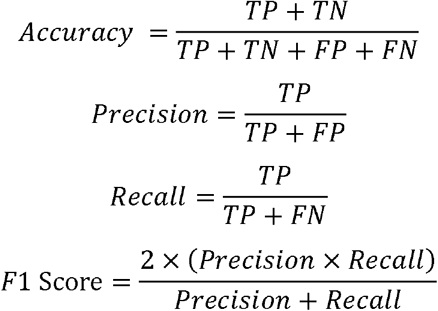

To further assess model stability and reliability standard deviation (SD) and 95% confidence intervals (CI) were calculated.^44^ Additionally, ROC with AUC analysis was performed to assess discriminative performance.^45^

#### 2.3.5. Cross validation of ML model with explainable AI (XAI)

Local interpretable model-agnostic explanations (LIME) provide a methodical approach for interpreting individual predictions of drug resistance.^46,47^ For a given tumor sample, LIME generates perturbations around the instance by introducing small variations in input features (e.g., gene amplification status, ecDNA burden, drug properties) while maintaining biologically relevant information. These perturbed inputs are evaluated by the trained ML model to obtain local predictions. The surrogate model assigns relevance scores to each feature, reflecting its contribution to the predicted drug resistance for that specific tumor. This localized interpretability enables identification of the ecDNA-associated genes and drug properties that drive resistance in individual samples, supporting mechanistic insights and potential clinical decision-making.^25^ By analyzing these local variations, LIME constructs a simple, interpretable surrogate model *g* that approximates the behavior of the complex ensemble model *f* near that sample. The surrogate model is designed to be both accurate and interpretable, and its optimization is expressed as:

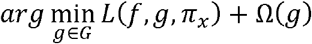

Here, *L* (*f,g, π*_*x*_) measures how well *g* replicates *f* in the neighborhood of sample *x, π*_*x*_ defines the locality or proximity of perturbed instances to the original tumor, and Ω (*g*) penalizes model complexity to ensure interpretability.

Another XAI method used in this study is Shapley Additive Explanations (SHAP), which decomposes the prediction for each individual sample into additive contributions from its constituent features.^48,49^ In this framework, model outputs are expressed as the sum of feature-specific Shapley values, enabling quantitative interpretation of how ecDNA-related genomic features and drug physicochemical properties influence resistance predictions. The Shapley value is formally defined as:

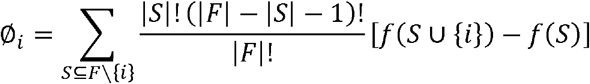

Here, *F* is the set of all features, *S* is a subset excluding feature *i*, and *f*(*S*) epresents the model prediction based on feature subset *S*. This formulation ensures that the contribution of each feature is fairly quantified, considering all possible feature interactions.

#### 2.3.6. Clinical simulation of gene specific chemotherapy resistance in TNBC

To further evaluate the potential clinical applicability of the trained Voting Ensemble model, a simulation framework was implemented to estimate drug resistance probabilities across ecDNA-associated genes expressed in the 4T1 TNBC model (*Fig. 3, Table 1*). The predictive features, including ecDNA burden, ecDNA prevalence, and drug LogP, were selected based on interpretability analyses using LIME and SHAP. Three chemotherapeutic agents, such as DOX, PAC, and Hydroxyurea (HU), were evaluated for chemotherapy resistance. The model predicted gene-specific resistance probabilities for each therapeutic agent and a resistance threshold of 0.60 was applied to classify high-risk resistance patterns.

**Table 1.**
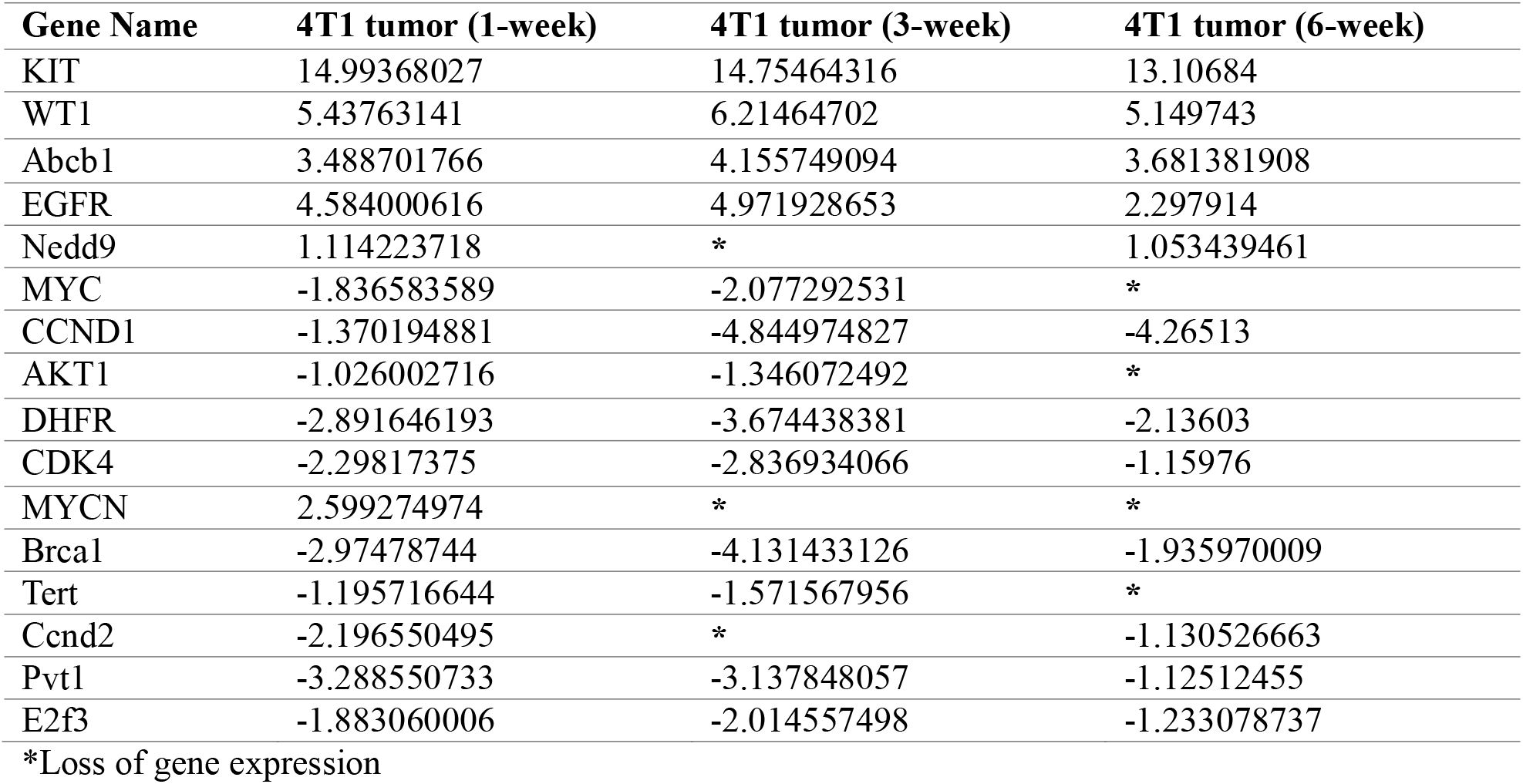
Gene expression level of selected ecDNA genes across different stages of tumor growth (Log2FC)

## 3. Results

### 3.1. Transcriptional shift in 4T1 cells: Insights from DEG analysis

Characterizing global transcriptional remodeling across distinct time points during the course of tumor progression could provide essential context for interpreting the temporal behavior of genes previously reported to be associated with ecDNA. Because ecDNA-associated genes function within the broader transcriptional network of the tumor,^50^ their stage-specific expression patterns could be interpreted relative to overall gene expression changes. DEG analysis provides a quantitative framework to capture the extent of transcriptional reprogramming across defined stages of tumor growth.^51^ As indicated in *Fig. 1A-C*, during the early time point (1w) in an *in vivo* model system, 4T1 tumor cells showed substantial transcriptional differences relative to the baseline 4T1 cell line, as 14,571 genes were differentially expressed (which is similar to the previous report^52^), with 6,507 genes upregulated. At the intermediate stage (3w), transcriptional differences remained prominent, with 6,799 genes upregulated. By the late stage (6w), 4T1 tumor samples showed a further transcriptionally diverged state, with only 8,925 gene remaining differentially expressed while 3,973 genes were upregulated. These results indicate that roughly 60% of the genes differentially expressed at 1w were no longer differentially expressed at 6w, suggesting cumulative stage-associated transcriptional remodeling and/or shifts in tumor-state composition.

**Fig. 1:**
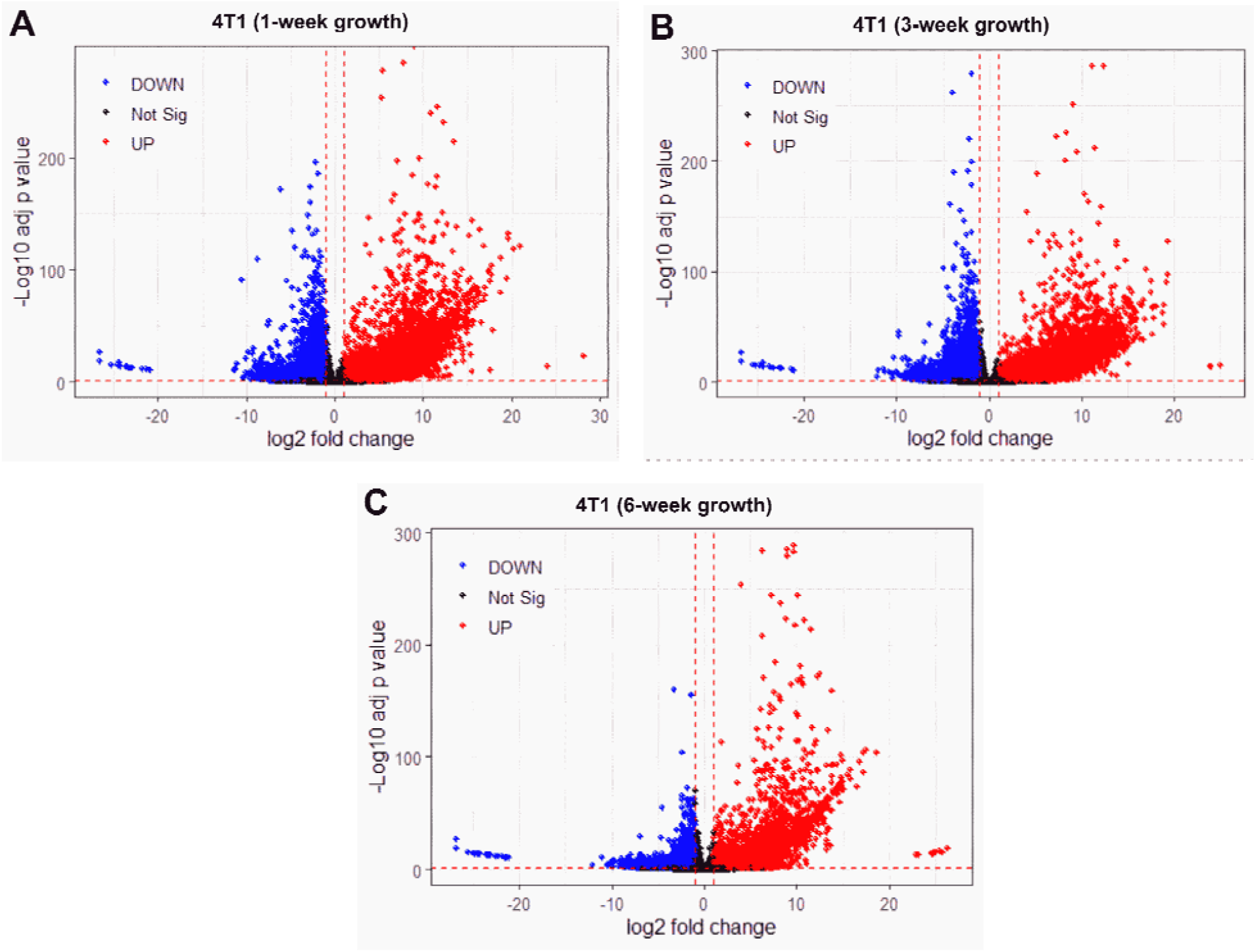
Volcano plot at different stages of 4T1 tumor growth. Using 4T1 cell RNA-seq as a baseline, thes plots show gene expression changes at (A) 1-week (14,571 DEGs), (B) 3-week (15,036 DEGs), and (C) 6-week (8,925 DEGs). The x-axis (log2 fold change) indicates the magnitude and direction of expression changes, with positive values denoting higher expression and negative values denoting lower expression.

The y-axis (labeled as “Log10 adj. p-value”) reflects statistical significance, where higher values (e.g., 0) correspond to lowest adjusted p-values (e.g., 1e-200), highlighting genes with highly confident differential expression. The plot reveals numerous genes with large fold changes (far left/right on the x-axis) and extreme significance (top of the y-axis), suggesting substantial molecular shifts during tumor progression. DOWN - downregulation; Not Sig-not significant; UP - upregulation.

Principal Component Analysis (PCA) also highlights transcriptional separation between the baseline 4T1 cell line and tumor samples across time points (*Fig. 2*). PCA provides a low-dimensional representation of global gene expression differences across groups. For instance, 4T1 cell line (red) was separated from all tumor samples along Principal Component 1 (PC1; 81% variance), indicating a major transcriptomic shift between *in vitro* baseline and *in vivo* tumor samples. The tight clustering of 1w and 3w groups indicates similarities in gene expression patterns across these time points. In contrast, 6w samples showed greater separation from earlier time points, consistent with higher divergence of transcriptomic remodeling at later stages (*Fig. 2*).^53^ Consistent separation of 6w samples in both DEG counts (*Fig. 1*) and PCA spac (*Fig. 2*) supports the hypothesis that later time points show a more distinct global transcriptional remodeling relative to earlier stages.

**Fig. 2:**
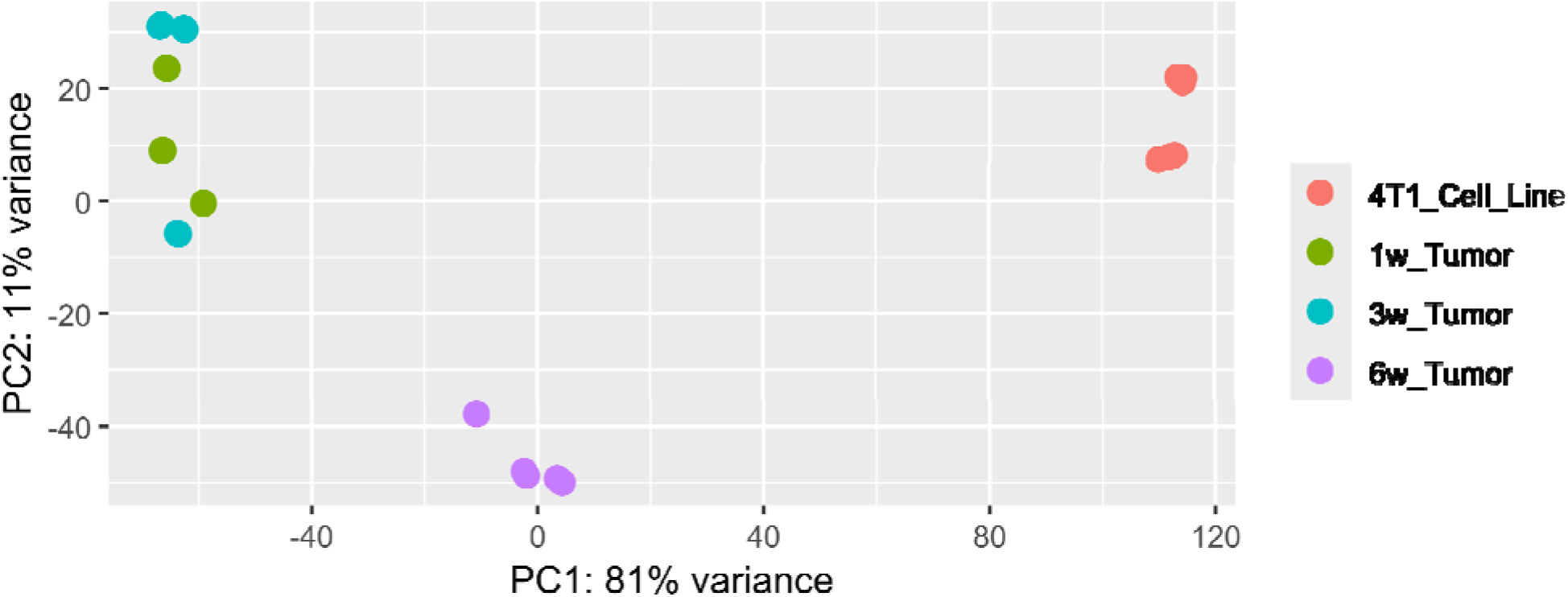
Principal Component Analysis (PCA) for 4T1 cells (baseline) and 1w, 3w, and 6w samples.

### 3.2. Temporal molecular remodeling of ecDNA-related genes

EcDNA has been reported to be associated with oncogene amplification and altered transcriptional output in multiple cancer contexts.^54^ Thus, after establishing global transcriptional differences across time point (*Figs. 1 and 2*), we next asked whether genes previously reported to be ecDNA-associated show stage-associated expression patterns. We identified 16 genes (out of 33 listed ecDNA gene observations; *se Supplementary Data Table*), which were reported by others as ecDNA-associated,^2,55-60^ expressed in 4T1 samples (*Fig. 3, Table 1*). Similar to the global gene expression patterns (*Figs. 1 and 2*), th transcriptional landscape of ecDNA-related oncogenes also showed dynamic shifts between weeks 1 to 6. For example, early and intermediate stages (1w and 3w) shared similar gene expression patterns, wherea by late-stage (6w), a major transcriptional reprogramming occurred, indicated by the downregulation or loss of certain ecDNA genes (*Fig. 3, Table 1*).

For instance, *MYC*, a well-known ecDNA oncogene,^2^ was highly expressed during the early time point but was progressively downregulated and eventually lost by 6w. This shift was accompanied by increased expression of *WT1*, another ecDNA-associated oncogene.^55^ Additionally, the expression of the ecDNA-related gene, *EGFR*,^2^ was decreased by approximately 50% from 1w to 6w, indicating a potential stage-associated reduction in EGFR signaling. In contrast, the well-known PAC resistance gene, *ABCB1*,^61^ remained consistently expressed across different stages. Meanwhile, *NEDD9*, a key regulator of cell migration and invasion,^2^ followed a distinct transcriptional pattern: it was initially expressed at 1w, suppressed at the intermediate stage (3w), and re-emerged by 6w (*Fig. 3, Table 1*). Collectively, these expression patterns indicate timepoint-dependent remodeling of ecDNA-associated genes.

Because prior studies have reported associations between tumor growth dynamics and sequence-level variation in ecDNA-related loci,^19^ we asked whether transcript-derived sequence differences could be observed across time points in selected ecDNA-related genes. We examined aligned RNA-seq reads for selected ecDNA-related genes (e.g., *CDK4* and *EGFR*) across three time points. We observed increasing numbers of nucleotide differences relative to the reference sequence. For instance, in *CDK4* and *EGFR*, the number of nucleotide differences increased from ~374 (1w) to 607 (3w) to 749 (6w) and from ~3 (1w) to 15 (3w) to 228 (6w), respectively. A representative region of *CDK4* with its sequence-difference patterns across the three stages is shown in *Fig. 4*. These observations indicate timepoint-associated increases in transcript-derived sequence differences in the examined regions; however, RNA-seq-based differences do not establish genomic mutations and should be interpreted as putative transcript-level variation.

**Fig. 3:**
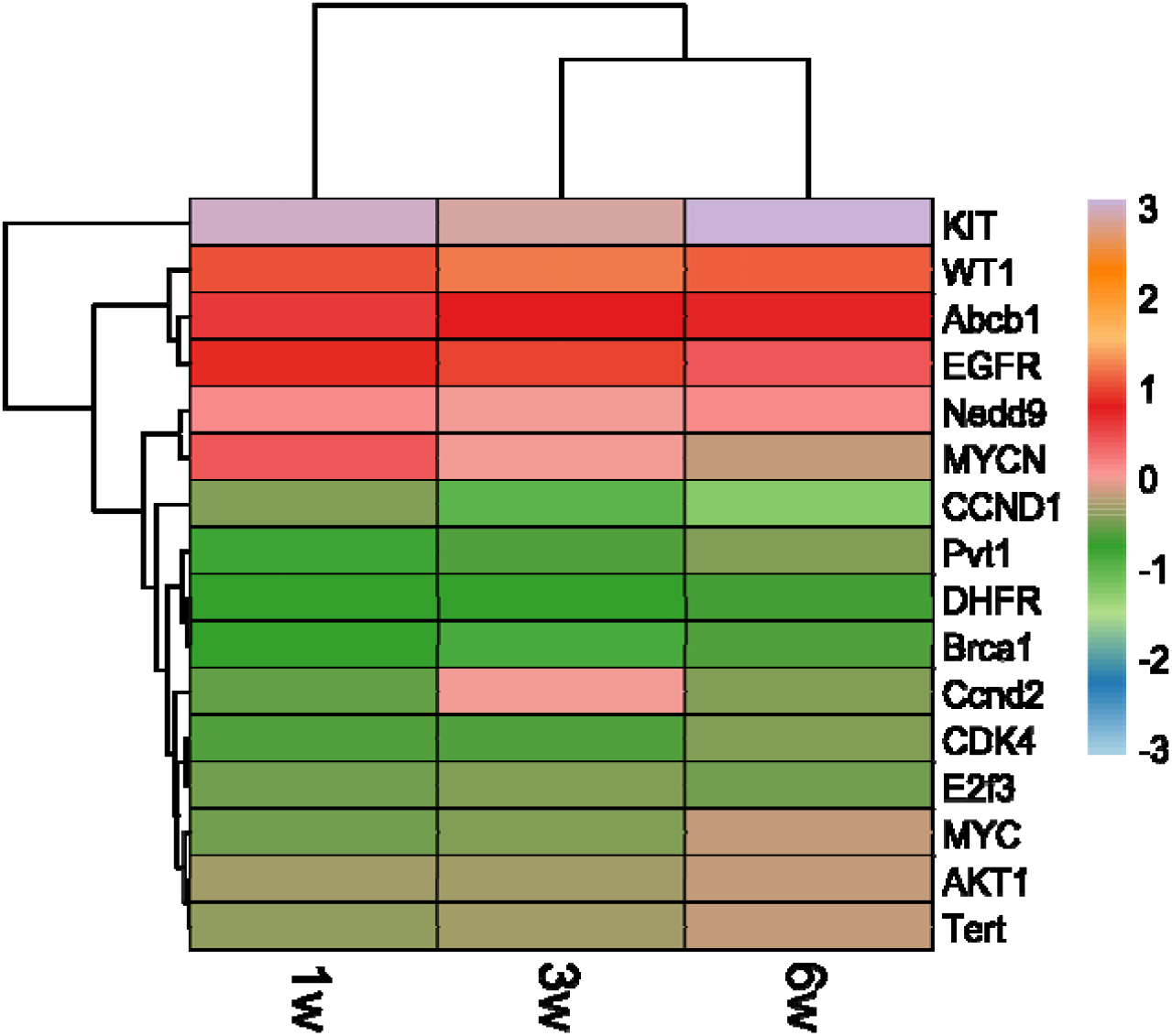
Changes in expression for ecDNA-associated genes across different stages of tumor development (1-, 3-, and 6-week post-tumor-implantation).

**Fig. 4:**
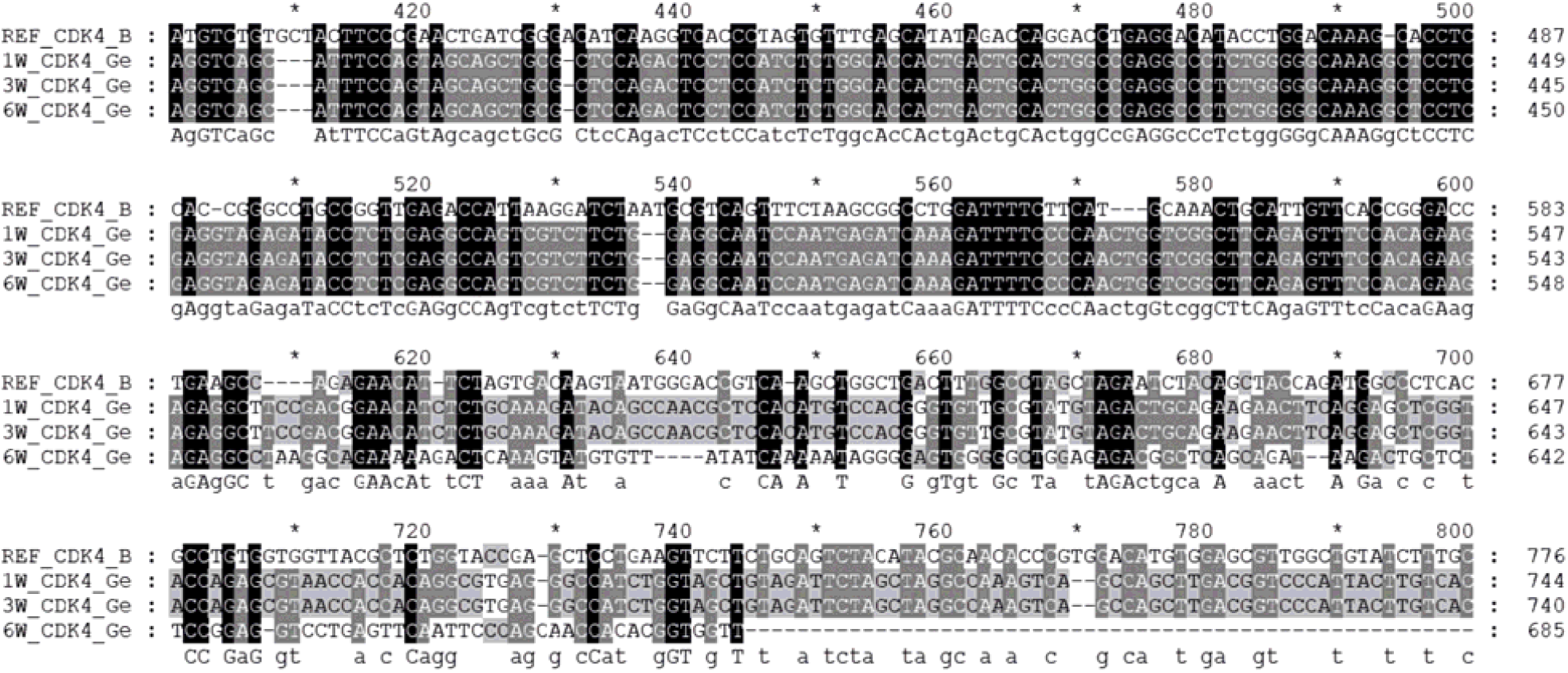
Sequence alignment of the CDK4 gene across experimental groups. We randomly extracted the CDK4 gene sequence from one of the six replicates from each time point: 1-week (1w), ID#SRR10428183; 3-week (3w), ID#SRR10428189; 6-week (6w), ID#SRR10428195; and aligned them with the Mus musculus reference CDK4 sequence (GenBank Accession ID #BC031599). Across the aligned region, we detected approximately 374 (1w), 607 (3w), and 749 (6w) nucleotide variations relative to the reference sequence. Among these, the number of nucleotide deletions was 21 (1w), 30 (3w), and 36 (6w), while insertions were 1 (1w), 16 (3w), and 21 (6w). For better visualization and due to space constraints, a representative segment of the alignment is shown, highlighting the regions with the highest mutation density.

### 3.3. Interaction between ecDNA-related proteins and anti-TNBC drugs

Because ecDNA-associated oncogenes have been linked to resistance phenotypes,^2,19^ transcript-derived sequence variation in ecDNA-related genes could plausibly correspond to altered predicted drug-protein interactions, which we explored using *in silico* modeling. Although we cannot fully predict all ecDNA gene(s) that are likely to be responsible for ecDNA-driven drug resistance, we selected the ecDNA-associated oncogene *ABCB1* (P-glycoprotein or Multidrug Resistance Protein 1) to investigate predicted interactions with DOX and PAC. *ABCB1* is an established mediator of chemotherapy resistance in cancer,^2,55-60,62-67^ including in TNBC,^64^ highly expressed across three stages (*Fig. 3, Table 1*). Molecular docking analysis provides a hypothesis-generating approach to estimate relative binding affinity differences between modeled protein variants and drugs.^62,63^ Here, we evaluated whether wild-type versus variant protein structures differ in predicted drug-protein binding interactions.

DOX interacts with ABCB1.^56^ The molecular docking analysis showed that wild-type ABCB1 strongly binds DOX (−10.5 kcal/mol), stabilized by hydrogen bonds with Gln-725 and Ser-979 (*Fig. 5A*). In contrast, ABCB1 mutation leads to alteration in the binding pocket, forming new hydrogen bonds (Ala-1111, Arg-1192, and Gln-1193) but showing a reduced docking score (−7.5 kcal/mol) (*Fig. 5B*), suggesting weakened affinity and potential resistance. Similarly, wild-type ABCB1 showed strong binding to PAC (−16.8 kcal/mol), interacting with Gln-725 (*Fig. 5C*). In contrast, the mutant ABCB1 displayed a reduced binding affinity (−12.0 kcal/mol), attributed to the loss of hydrogen bonds and the emergence of weaker van der Waals interactions (Ile-190, Ile-352, and Phe-40) (*Fig. 5D*), consistent with mutation-driven conformational changes and potential resistance.

**Fig. 5:**
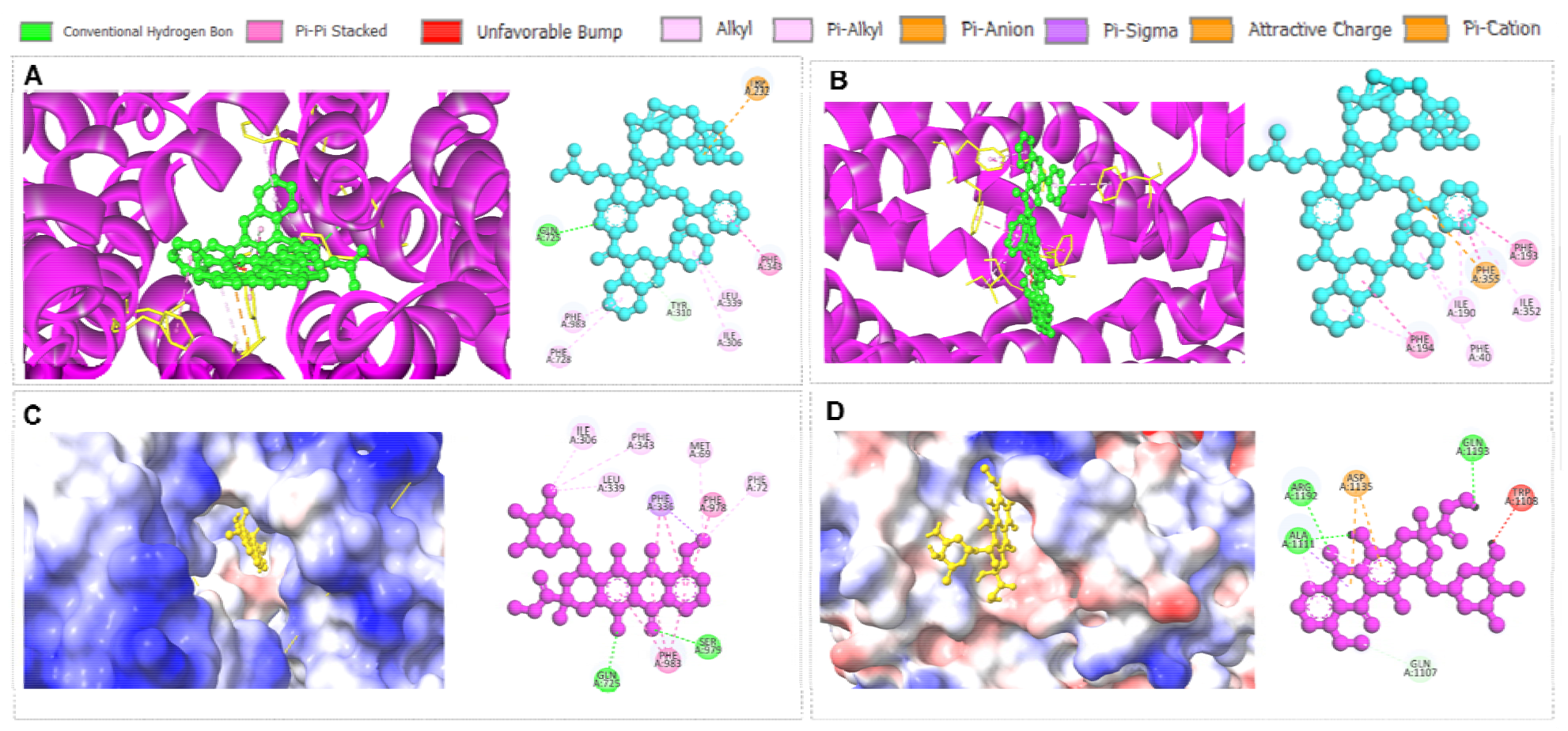
**(A-B)** Doxorubicin interaction with wild-type and mutant ecDNA proteins. (A-B) Doxorubicin interaction with (A) wild-type and (B) mutant ABCB1 protein. **(C-D)** Paclitaxel interaction with wild-type and mutant ecDNA proteins. (C-D) Paclitaxel interaction with (C) wild-type and (D) mutant KIT protein, whereas mutant structures were derived from 4T1 samples to compare potential differences in predicted drug–protein binding interactions.

### 3.4. EcDNA burden stratifies drug resistance

To complement transcript-level analyses with a broader resistance-focused perspective, we next evaluated whether ecDNA-related features derived from a curated literature dataset are associated with resistanc classification in a predictive modeling framework. Following the expansion of the final dataset into 1,160 data points as outlined in the Methodology section, initial exploratory analysis was performed to understand the distribution of the target variable and its relationship with crucial biological and pharmacological features. The drug resistance distribution showed a moderate class imbalance, with resistant samples (n ~ 840) exceeding sensitive samples (n ~ 320) (*Fig. 6A*). The ecDNA Burden Score showed a strong positive association with resistance probability, with a near-monotonic increase in resistance as burden levels rose from 1 to 3. Samples with moderate to high ecDNA burden (scores 2-3) were predominantly resistant and had similar probability of resistance, whereas those with less or no detectable ecDNA burden (score 1) showed minimal probability of resistance (*Fig. 6B*). Additionally, comparison of drug molecular weight distributions (*see Supplementary Data Table*) across analyzed datasets showed that sensitive cohorts showed a pronounced and isolated peak at lower molecular weights (<200 g/mol), highlighting that smaller molecules encountered fewer resistance barriers (*Fig. 6C*). In contrast, the resistance cohort was shifted toward higher molecular weights, with a dominant peak around 450-500 g/mol and a secondary peak near 850 g/mol, indicating greater heterogeneity and a tendenc toward larger compounds. Notably, the near absence of sensitive compounds beyond 600 g/mol supports the biological premise that high-molecular-weight drugs are more prone to resistance mechanisms including reduced membrane permeability and increased efflux transporter activity.^68–70^ To further evaluate class separability (resistant and sensitive) within the multidimensional feature space, LDA wa performed, demonstrating robust separation between resistant and sensitive groups compared to non-LDA-optimized datasets (*Fig. 7*). Overall, our ML-based pre-processing strategy allowed these datasets to fit into further ML-based model training for prediction of ecDNA-driven chemotherapy resistance.

**Fig. 6:**
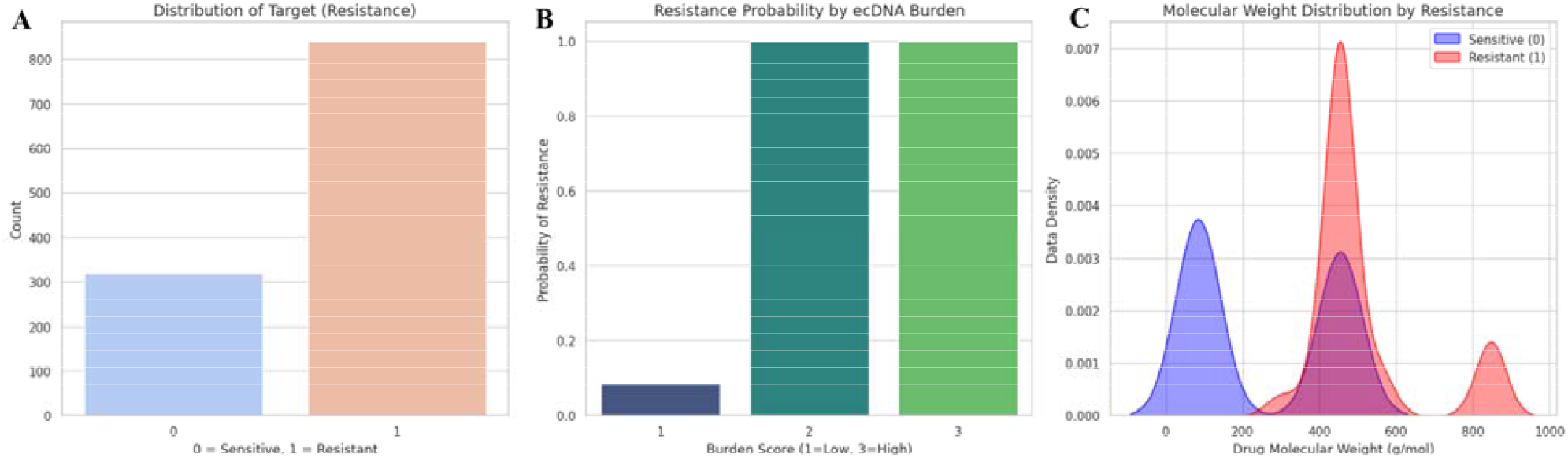
Correlation of ecDNA burden and drug resistance: **(A)** Distribution of resistant and susceptibl samples in the dataset, **(B)** Resistance probability across ecDNA burden scores, **(C)** Drug molecular weight distribution across datasets.

**Fig. 7:**
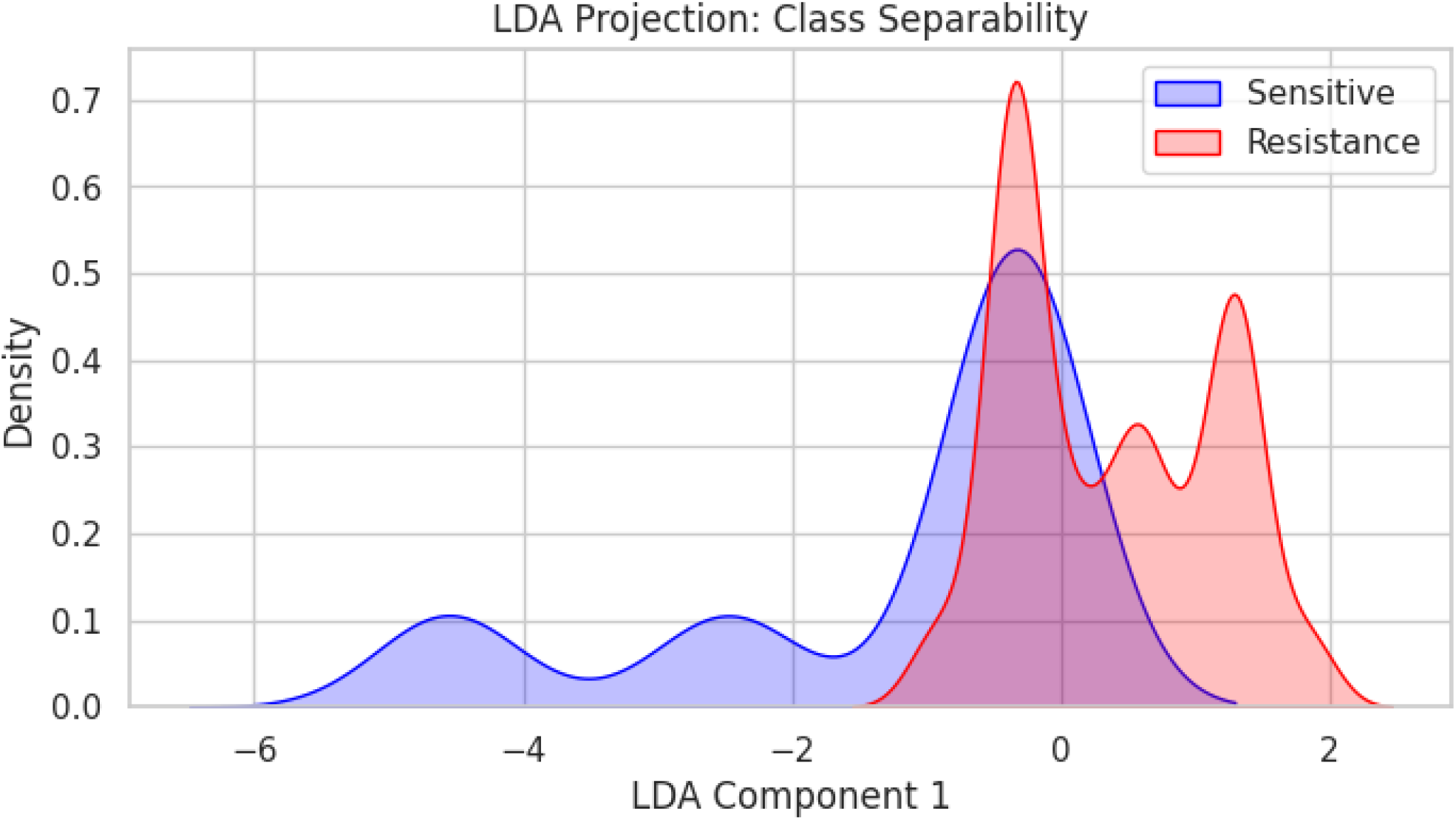
Feature based separation of ecDNA associated resistance states

### 3.5. Non-linear models capture ecDNA associated resistance signal

We next tested whether non-linear classifiers capture resistance-associated patterns in the ecDNA-feature dataset and compared their performance with a linear baseline model. The evaluated ML models, such a RF and XGBoost, achieved 100% performance in all criteria evaluated including accuracy, precision, recall, F1-score, and specificity (**Table 2**). However, these superior performances were likely due to overfitting caused by the lack of larger experimental datasets^71^. In contrast, the Logistic Regression model achieved 88.12% accuracy and an F1-score of 88.03%. While precision and specificity remained at 100%, recall decreased to 78.62%, indicating a higher false-negative rate compared to other three models (**Table 2**). We then employed the Voting Ensemble model that combines all individual models to improve predictive accuracy, reduce overfitting, and better capture complex, non-linear relationships in the datasets compared to using an individual model alone.^72^ However, the Voting Ensemble model also produced 100% performances across all five tested criteria, and this was likely due to overfitting associated with the inclusion of small experimental datasets.^71^ Despite knowing this fact, we utilized the trained Voting Ensemble model for the post-processing ML prediction analysis.

**Table 2.** Performance of different ML models.

| Model | Accuracy | Precision | Recall | F1 Score | Specificity |
| --- | --- | --- | --- | --- | --- |
| Random Forest | 100.00 % | 100.00 % | 100.00 % | 100.00 % | 100.00 % |
| XGBoost | 100.00 % | 100.00 % | 100.00 % | 100.00 % | 100.00 % |
| Logistic Regression | 88.12 % | 100.00 % | 78.62 % | 88.03 % | 100.00 % |
| Voting Ensemble | 100.00 % | 100.00 % | 100.00 % | 100.00 % | 100.00 % |

We then further cross-validated the trained the Voting Ensemble model. The confusion matrix of the Voting Ensemble model demonstrated 116 true negatives (sensitive correctly predicted), and 145 true positives (resistant correctly predicted) (*Fig. 8A*). ROC analysis showed discriminative ability with an AUC of 1.00 (*Fig. 8B*), suggesting it can distinguish resistant and sensitive samples with up to 100% certainty based on the selected features (e.g., molecular weight). Learning-curve analysis showed that training accuracy increased from ~0.98 to 1.00 with increasing sample size, while cross-validation accuracy improved from ~0.25 to approximately 0.97-0.98 and converged with the training curve (*Fig. 8C*). Log loss decreased with increasing sample size, with cross-validation loss declining from ~1.47 to ~0.24. The final ensemble achieved a test-set log loss of 0.1225 (*Fig. 8D*). Because the long-format reshaping increases observation counts by representing multiple gene entries per sample, these performance metrics should be interpreted in the context of the dataset structure and should not be taken as evidence of generalizability.

**Fig. 8:**
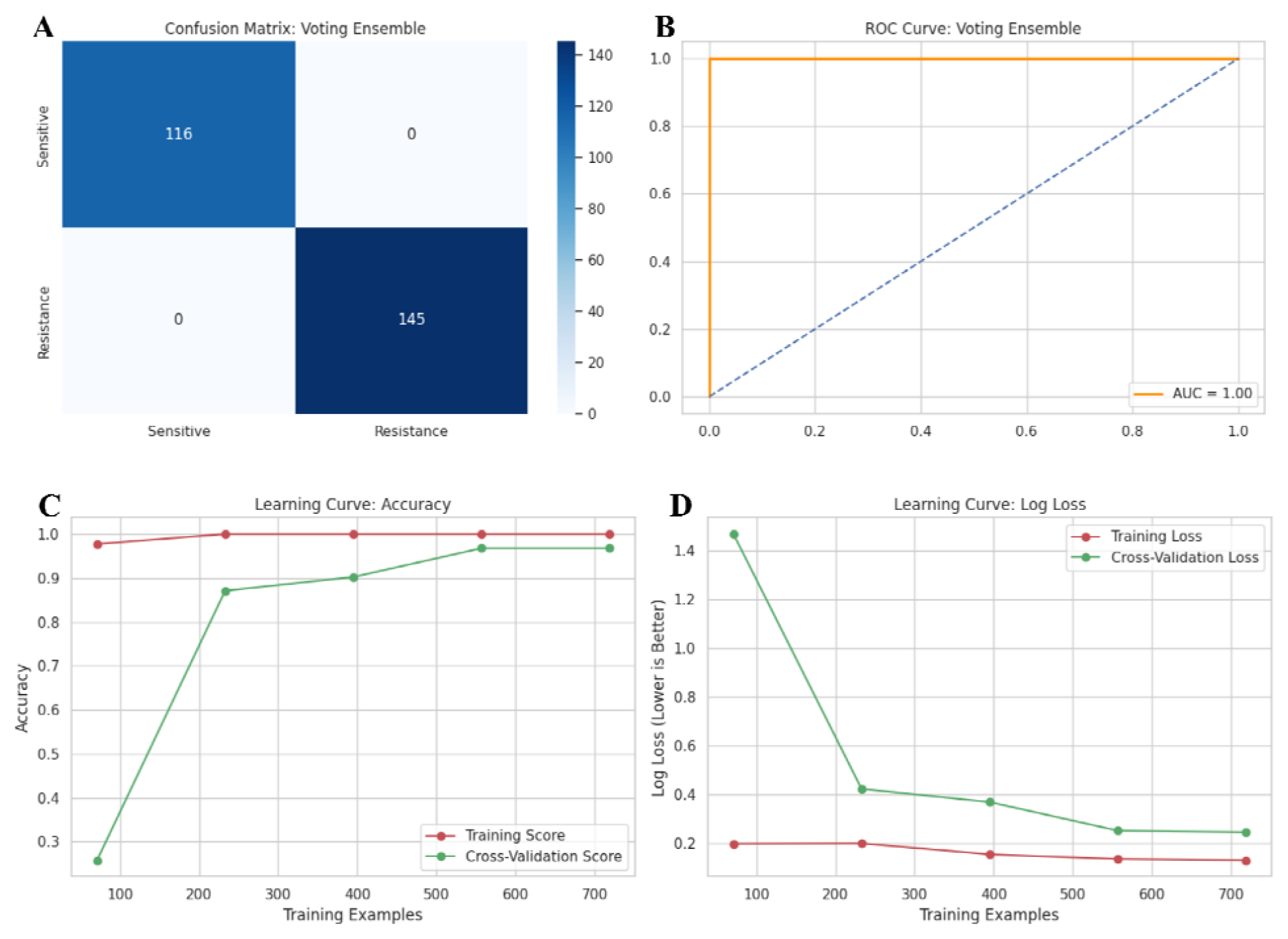
Performance evaluation of the model: **(A)** Confusion matrix for classification accuracy and **(B)** ROC curve for model discrimination. Learning curve analysis of model accuracy **(C)** and log loss **(D)**.

To further assess model confidence and probabilistic separation, the distribution of predicted resistance probabilities was examined on the independent test set using the Kernel density estimation (KDE). The Voting Ensemble’s predicted probabilities for sensitive samples (*y = 0*) were concentrated near 0.0, whereas probabilities for resistant samples (*y = 1*) clustered near 1.0, showing a clear bimodal separation. More specifically, sensitive samples were tightly clustered within a narrow probability range of ~0.10-0.20, indicating consistently low predicted resistance probabilities. In contrast, resistant samples were distributed predominantly between approximately 0.75 and 1.00, with density peaks near ~0.80 and ~0.95 (*Fig. 9*). The bimodal probability distribution indicates that the Voting Ensemble model assign consistently low resistance probabilities to sensitive samples and high resistance probabilities to resistant samples, demonstrating strong probabilistic separation between classes within this dataset (*Fig. 9*). This separation suggests that the combined feature set used in the model contains substantial information relevant to resistance classification, enabling the Ensemble model to distinguish resistant and sensitive profiles with high confidence within the dataset. Importantly, because the long-format dataset structure increases the number of observations by representing multiple gene entries per sample, the resulting probability separation reflects patterns present within the dataset rather than validated predictiv performance in independent cohorts. Therefore, these findings indicate that the curated feature set contains a strong resistance-associated signal within the dataset, providing a basis for hypothesi generation and motivating further analyses to identify which biological features most strongly drive model predictions.

**Fig. 9:**
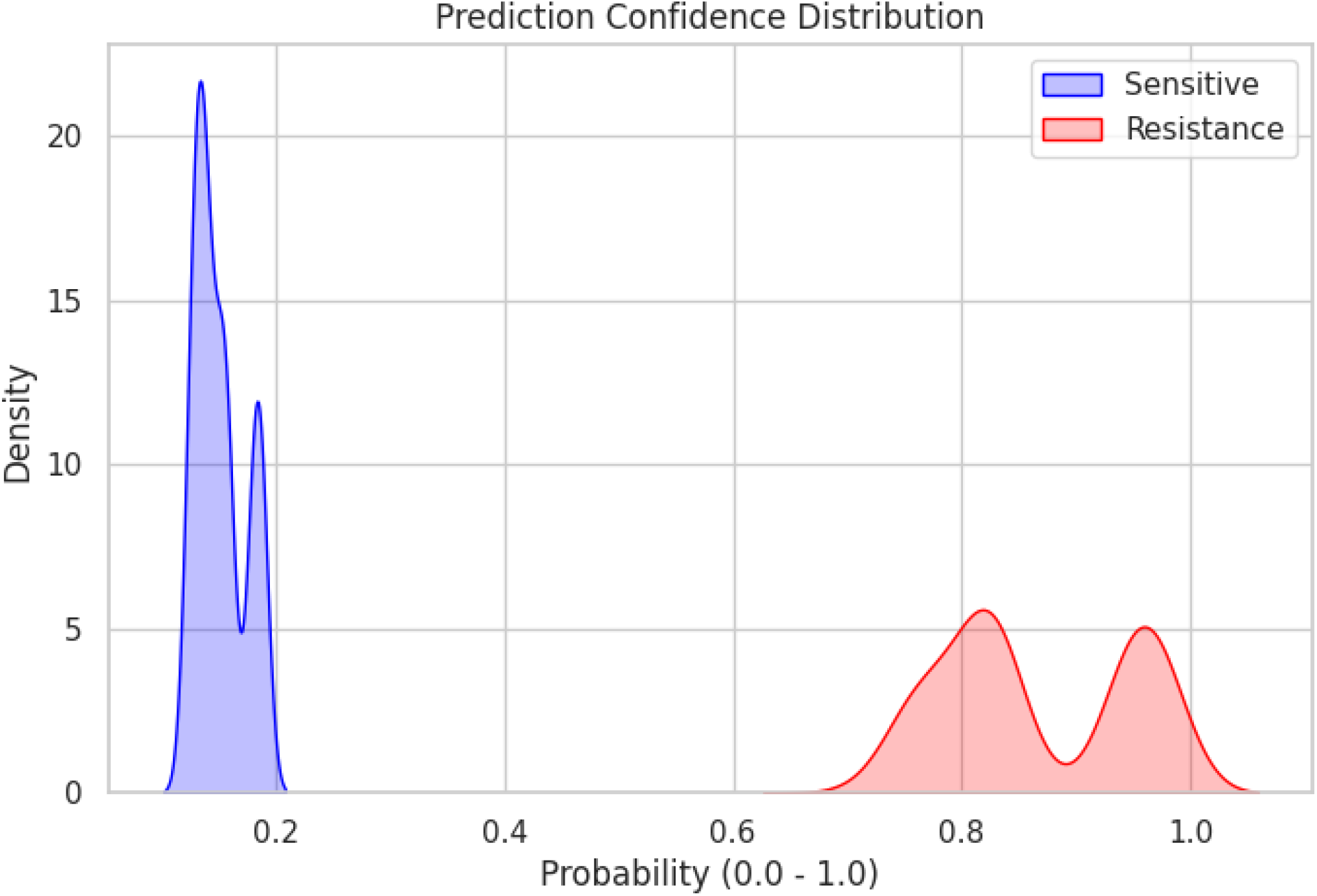
Kernel density distribution of ecDNA associated predicted resistance and sensitive probabilities.

### 3.6. Explainability AI (XAI) reveals ecDNA as a dominant determinant

Because high model performance can mask the “black-box” nature of the decision logic, we applied explainability methods to identify which ecDNA- and drug-related features contributed most to resistance classification. XAI refers to a set of methods and tools designed to make ML models understandable to humans, bridging the gap between complex “black-box” algorithms and clinical decision-making.^73^ Thus, to complement model evaluation, XAI approaches were applied to ensure transparency and interpretability of predictions. The SHAP beeswarm plot, which provides a global view of model behavior by illustrating the distribution and magnitude of feature contributions across all samples in the dataset,^74^ demonstrates that ecDNA burden score and ecDNA prevalence score exert the greatest impact on model output. High ecDNA-feature values (red) were predominantly associated with positive SHAP value (shifting predictions toward resistance), whereas lower ecDNA feature values (blue) were associated with negative SHAP contributions (shifting predictions toward susceptibility) (*Fig. 10*). Within this predictive framework, ecDNA-related metrics emerged as dominant contributors to resistance classification. Drug-related features such as Drug LogP and Drug Molecular Weight also contributed to model predictions, but with comparatively smaller magnitudes (*Fig. 10*).

**Fig. 10:**
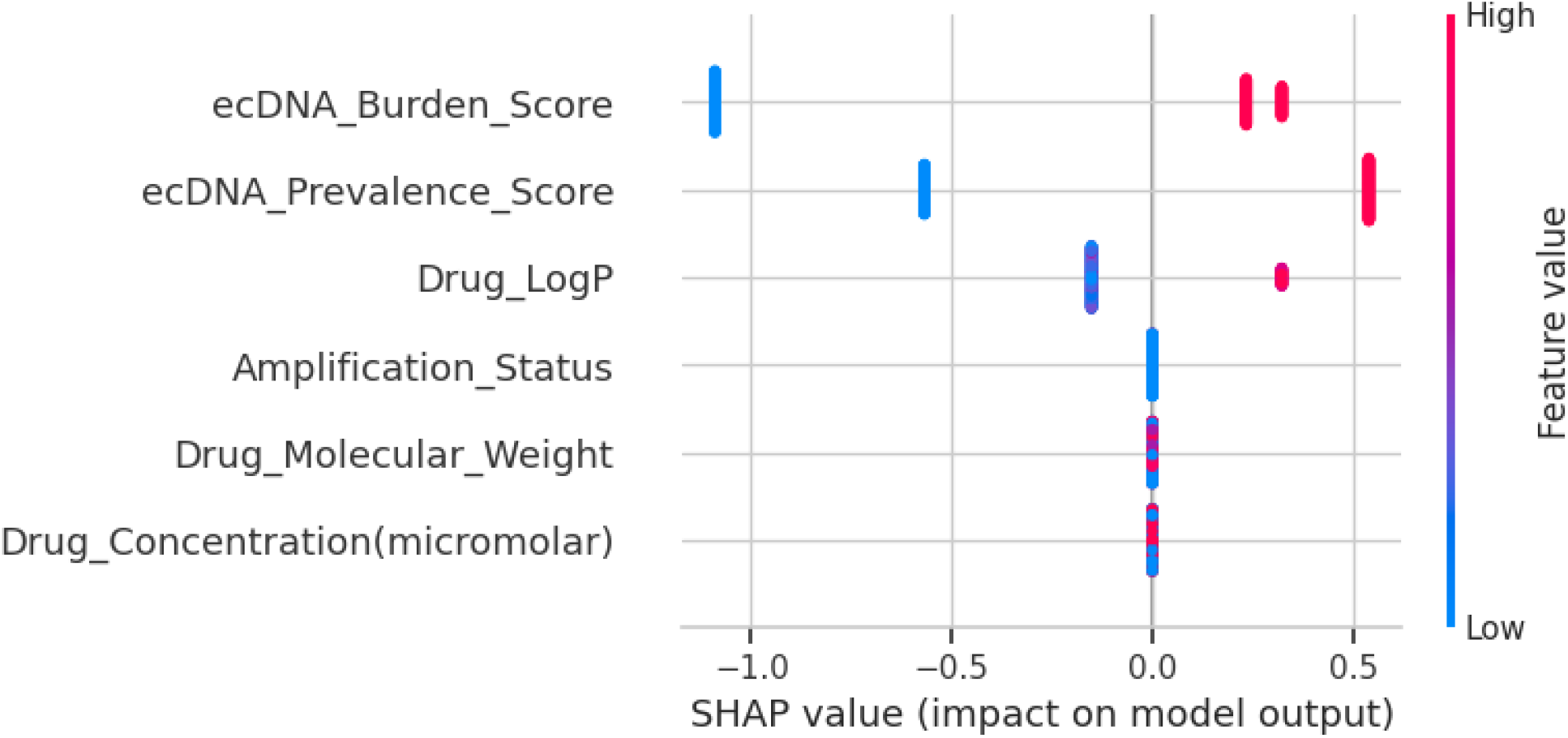
SHAP values of several important features contributions to resistance prediction.

To further complement SHAP interpretation, LIME was applied to interpret an individual prediction to allow identification of features that most strongly influence a specific prediction.^48^ The LIME model predicted a resistance probability of 82%, while probability of sensitivity was 18%. The LIME model indicated that elevated ecDNA Prevalence Score (1.21) and ecDNA Burden Score (1.34) contributed most strongly to the resistant prediction (*Fig. 11*). These results are consistent with the global SHAP interpretation in which higher ecDNA metrics align with resistance classification within the model.

**Fig. 11:**
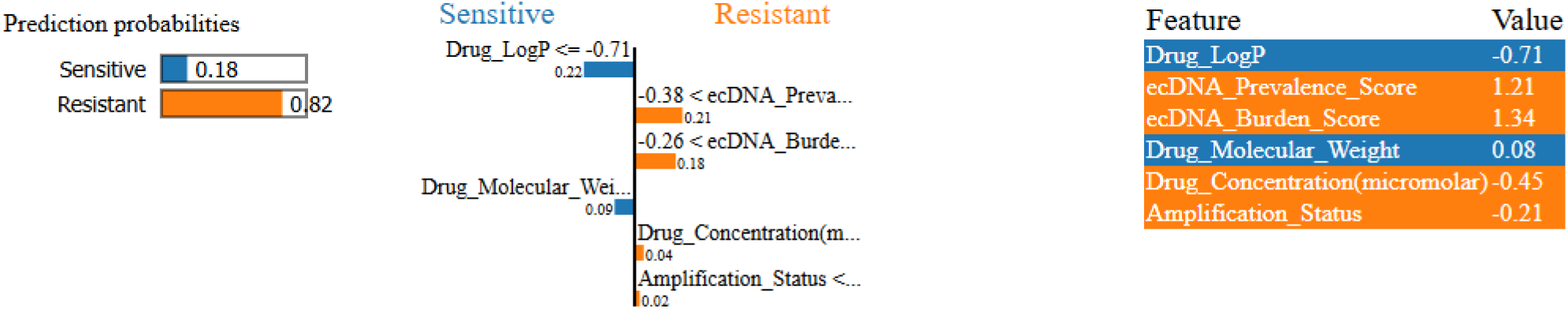
Lime based percentage attribution of predictive features in resistance modeling.

### 3.7. Drug response profiling of ecDNA in TNBC

After observing that ecDNA-associated features were strongly associated with resistance classification in the curated dataset, we extended the analysis using an ML-based simulation to explore how PAC and DOX might behave in an ecDNA-feature context. This analysis was conducted across 16 ecDNA-related genes (*Fig. 3, Table 1*) for which direct experimental validation of PAC and DOX responses was not available. Resistance probabilities were computed while maintaining high ecDNA burden and systematically varying amplified gene identity. The resulting predictions indicated elevated predicted resistance risk for PAC (95%) across the gene panel, whereas DOX showed comparatively moderate predicted resistance (78%) (*Fig. 12*). Importantly, the relative uniformity of resistance predictions acros multiple amplified oncogenes suggests that, under conditions of high ecDNA burden, drug physicochemical characteristics may exert a greater influence on predicted therapeutic response than individual gene amplification status alone.

**Fig. 12:**
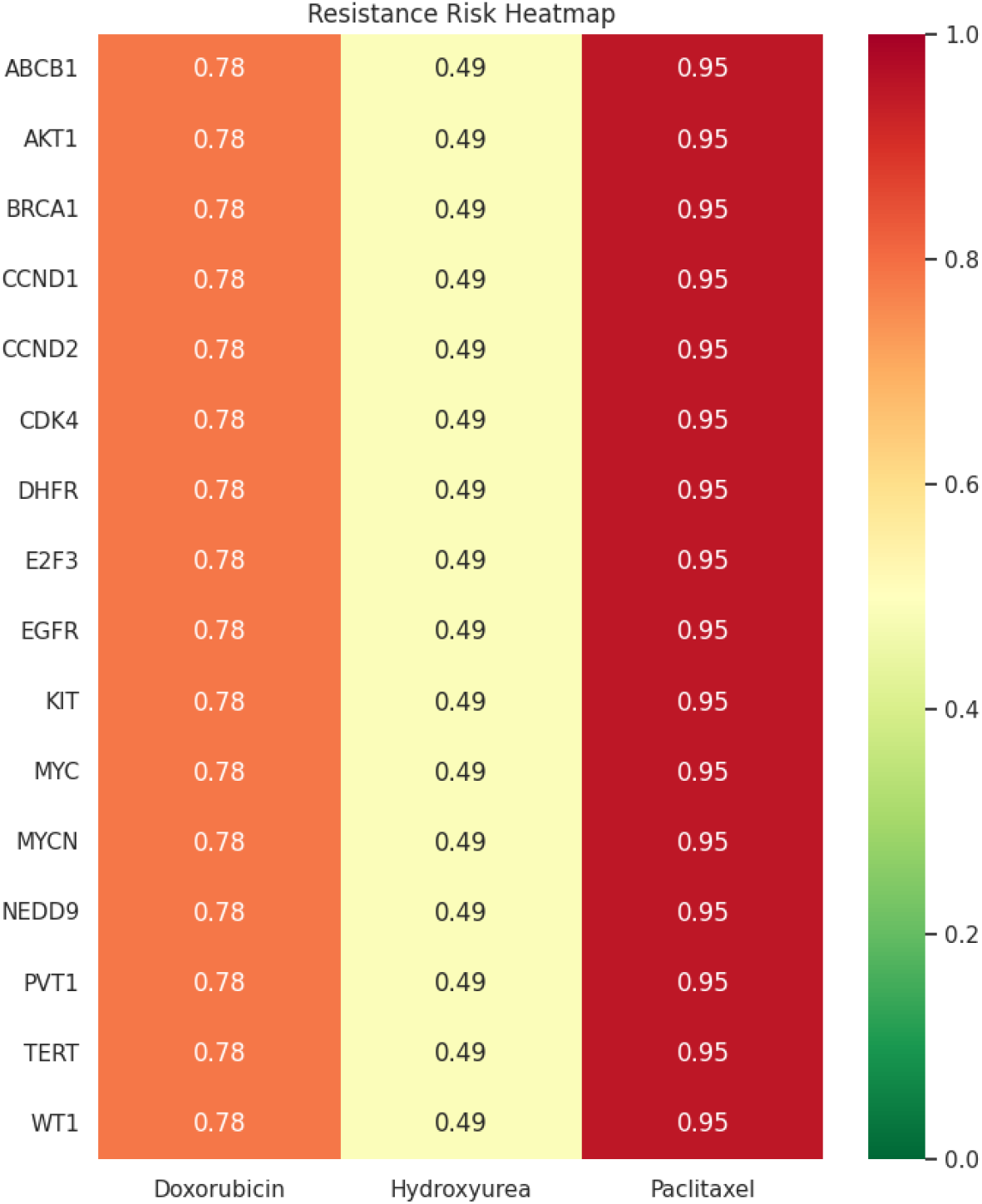
Predicted resistance risk across ecDNA genes in TNBC. Color scale represents predicted resistance probability (0 = sensitive, 1 = resistant), with green indicating low resistance and red indicating high resistance.

Hydroxyurea (HU), which inhibits ribonucleotide reductase, has been reported to induce replication stress and to reduce ecDNA content in certain experimental systems.^75^ Since HU can target and deplete ecDNA, we incorporated HU into the resistance simulation an additional drug context. Although previous studies indicate that replication-stress-inducing drugs may preferentially affect ecDNA-containing tumor cells, a systematic evaluation of HU sensitivity across ecDNA-amplified gene contexts remains limited. The SHAP and LIME features that were used to evaluate predicted resistance for DOX or PAC were incorporated in HU simulation. HU showed lower predicted resistance probabilities compared with PAC and DOX across the 16 ecDNA-associated genes (*Fig. 3, Table 1*), with the predicted resistance risk for HU was 49%, which was ~1.6- and 1.9-fold lower than DOX and PAC, respectively, suggesting that HU-driven ecDNA depletion enhanced predicted sensitivity to DOX or PAC *(Fig. 12)*.

## 4. Discussion

Chemotherapy resistance remains a major challenge in the treatment of many cancers, including TNBC. Recently, ecDNA has been linked as a potential contributor to chemoresistance in cancer.^2,19^ In this study, we applied an integrative computational framework combining transcriptomic profiling, structural modeling, and ML to explore the potential contribution of ecDNA-associated genes to chemoresistance in TNBC. Our global transcriptomic analyses revealed substantial transcriptional remodeling during tumor progression in the 4T1 TNBC model. For example, a relatively similar global expression pattern observed at 1w and 3w, whereas a more divergent transcriptional state observed at 6w, consistent with progressive molecular reprogramming. Likewise, for ecDNA-associated genes, similar expression patterns observed at early (1w) and intermediated (3w) stages, with a major transcriptional remodeling occurred at late stage, reflecting the global gene expression patterns. These findings suggest that ecDNA-associated genes may be temporally remodeled during tumor progression, and this is similar to what we reported previously for temporal molecular remodeling of T cells across various stages of 4T1 tumor growth.^6^ Beyond transcriptional dynamics, our exploratory analysis of transcript-derived sequence variation in selected ecDNA-associated genes such as *CDK4* and *EGFR* suggested increasing divergence from reference sequences across tumor stages. While RNA-seq-derived sequence differences cannot be interpreted as definitive genomic mutations without orthogonal validation,^75,76^ the observed patterns raise the possibility that tumor progression may be accompanied by increasing molecular heterogeneity, facilitating the emergence of tumor cell subpopulations with altered drug sensitivity.^77^ Indeed, intratumoral heterogeneity is a well-recognized hallmark of chemotherapy-resistant cancers and enables the selective survival of clones capable of tolerating therapeutic exposure.^78^

To further explore how ecDNA-associated variation might influence therapeutic response, we examined predicted interactions between selected ecDNA-related proteins and commonly used anti-TNBC drugs, such as anthracyclines (e.g., DOX) and taxanes (e.g., PAC).^16,17,79^ Both drugs can induce ecDNA amplifications, potentially influencing drug sensitivity or resistance.^62,63,80,81^ The docking analyses provide an additional layer of hypothesis generation by asking whether modeled variant proteins differ from wild-type proteins in predicted interactions with standard anti-TNBC drugs.^82^ Importantly, while wild-type ABCB1 protein showed moderate to strong binding affinity toward both DOX and PAC, the mutant ABCB1 showed a consistent reduction in binding affinity, indicative of a potential drug-resistant phenotype. More specifically, in case of DOX, although the mutant protein formed alternative hydrogen bonds (e.g., Ala-1111, Arg-1192, and Gln-1193), these interactions were likely not sufficient to maintain the stability observed in the wild-type complex, possibly due to mutation-induced alterations in the binding pocket. Similarly, the weak interaction of PAC with the mutant protein is likely associated with the loss of hydrogen bonds and/or the gain of less stable van der Waals interactions.^83,84^ These findings are consistent with previous reports indicating that structural and functional alterations in ABCB1 (P-glycoprotein) can significantly influence drug binding and efflux activity, ultimately contributing to multidrug resistance in cancer cells.^85,86^ Importantly, observations obtained from docking analysis strongly align with our ML findings. For instance, the predictive framework consistently identified ecDNA burden and prevalence as dominant determinants of drug resistance. Across cross-validation folds and independent test sets, resistance classification remained separable, with SHAP and LIME analyses confirming that elevated ecDNA metrics driving resistance predictions. This convergence between *in vivo* transcriptional remodeling and ML-based feature attribution supports the hypothesis that ecDNA associated genomic instability functions as a structural driver of therapeutic resistance rather than merely a correlated marker. Our ML-based clinical simulations in ecDNA in TNBC further demonstrated drug-specific resistance patterns. For example, PAC showed uniformly high predicted resistance (95%), whereas DOX demonstrated moderate resistance (78%) across 16 ecDNA genes. In contrast, HU, a replication stress inducing agent implicated in ecDNA dynamics,^87^ demonstrated comparatively lower predicted resistance (0.49), suggesting potential vulnerability of ecDNA-rich tumors to replication interference. Because ecDNA-containing cells rely on elevated replication and transcriptional activity, perturbing replication stress pathways may preferentially disrupt ecDNA maintenance and propagation.^87^

## 5. Conclusions

This study provides an integrative computational analysis involving temporal transcriptomic remodeling, modeled drug-target interactions, and predictive resistance modeling linked to ecDNA-associated genes. The findings suggest that ecDNA burden and prevalence are strongly associated with resistance patterns within the analyzed datasets. Although the results remain exploratory and require substantial experimental validation, our findings provide a coherent framework for investigating ecDNA as a contributor to chemotherapy resistance in TNBC. Future studies should comprehensively investigate the dynamics of ecDNA-associated genes using whole-genome sequencing (WGS) to characterize structural rearrangements, copy number alterations, and mutational trajectories over time and their potential role in chemoresistance. Additionally, investigations integrating longitudinal WGS, ecDNA structural mapping, functional drug response assays, and iterative ML modeling will be critical to determine how ecDNA could reshape resistance trajectories. Such integrative approaches may enable the development of ecDNA-informed predictive biomarkers and guide rational therapeutic strategies for cancers such as TNBC.

## Acknowledgments

D.S. was supported in part by a fund from the College of Science and Technology at North Carolina A&T State University (NCA&T), Faculty Fellowship Grant (GRADS-4C) at NCA&T, and by a grant from the NIH (1R16NS147983-01).

## Author Contributions

M.I.: Performed data analysis, prepared figures and tables, and wrote the manuscript; D.S.: Conceptualization, study outline, and supervision, and data analysis. All authors edited the manuscript and approved the submission.

## Data Availability

Data comprising sequence read counts and differentially expressed genes (DEGs) for the 1-, 3-, and 6-week periods are deposited under DOI: 10.5281/zenodo.16886936.

## References

1. Zhu H, Huangfu L, Chen J, Ji J, Xing X. Exploring the potential of extrachromosomal DNA as a novel oncogenic driver. Science China Life Sciences. 2025/01/01 2025;68(1):144–157. doi:10.1007/s11427-024-2710-3

2. Pecorino LT, Verhaak RGW, Henssen A, Mischel PS. Extrachromosomal DNA (ecDNA): an origin of tumor heterogeneity, genomic remodeling, and drug resistance. Biochem Soc Trans. Dec 16 2022;50(6):1911–1920. doi:10.1042/bst20221045

3. Huang Q, Zhang S, Wang G, Han J. Insight on ecDNA-mediated tumorigenesis and drug resistance. Heliyon. Mar 30 2024;10(6):e27733. doi:10.1016/j.heliyon.2024.e27733

4. Wu S, Bafna V, Chang HY, Mischel PS. Extrachromosomal DNA: An Emerging Hallmark in Human Cancer. Annual review of pathology. Jan 24 2022;17:367–386. doi:10.1146/annurev-pathmechdis-051821-114223

5. Wu S, Tao T, Zhang L, Zhu X, Zhou X. Extrachromosomal DNA (ecDNA): Unveiling its role in cancer progression and implications for early detection. Heliyon. 2023;9(11)doi:10.1016/j.heliyon.2023.e21327

6. Iftehimul M, Newman RH, Harrison SH, et al. Temporal molecular remodeling of T cells informs their possible adaptation in 4T1 tumors. Computational and Structural Biotechnology Journal. 2026/01/01/ 2026;31:169–178. doi:10.1016/j.csbj.2025.12.022

7. Yi E, Chamorro González R, Henssen AG, Verhaak RGW. Extrachromosomal DNA amplifications in cancer. Nature Reviews Genetics. 2022/12/01 2022;23(12):760–771. doi:10.1038/s41576-022-00521-5

8. Morales C, Ribas M, Aiza G, Peinado MA. Genetic determinants of methotrexate responsiveness and resistance in colon cancer cells. Oncogene. 2005/10/01 2005;24(45):6842–6847. doi:10.1038/sj.onc.1208834

9. Meng X, Qi X, Guo H, et al. Novel role for non-homologous end joining in the formation of double minutes in methotrexate-resistant colon cancer cells. Journal of Medical Genetics. 2015;52(2):135–144. doi:10.1136/jmedgenet-2014-102703

10. Nathanson DA, Gini B, Mottahedeh J, et al. Targeted Therapy Resistance Mediated by Dynamic Regulation of Extrachromosomal Mutant EGFR DNA. Science. 2014;343(6166):72–76. doi:doi:10.1126/science.1241328

11. Lange JT, Chen CY, Pichugin Y, et al. Principles of ecDNA random inheritance drive rapid genome change and therapy resistance in human cancers. bioRxiv. 2021:2021.06.11.447968. doi:10.1101/2021.06.11.447968

12. Pal Choudhuri S, Girard L, Lim JYS, et al. Acquired Cross-Resistance in Small Cell Lung Cancer due to Extrachromosomal DNA Amplification of MYC Paralogs. Cancer Discovery. 2024;14(5):804–827. doi:10.1158/2159-8290.cd-23-0656

13. Tyler LC, L. AT, Chen N, et al. MET gene amplification is a mechanism of resistance to entrectinib in ROS1+ NSCLC. Thoracic cancer. Nov 2022;13(21):3032–3041. doi:10.1111/1759-7714.14656

14. Morales C, García MJ, Ribas M, et al. Dihydrofolate reductase amplification and sensitization to methotrexate of methotrexate-resistant colon cancer cells. Molecular Cancer Therapeutics. 2009;8(2):424–432. doi:10.1158/1535-7163.mct-08-0759

15. Karami Fath M, Karimfar N, Fazlollahpour Naghibi A, et al. Revisiting characteristics of oncogenic extrachromosomal DNA as mobile enhancers on neuroblastoma and glioma cancers. Cancer Cell International. 2022/05/25 2022;22(1):200. doi:10.1186/s12935-022-02617-8

16. Smoots SG, Schreiber AR, Jackson MM, et al. Overcoming doxorubicin resistance in triple-negative breast cancer using the class I-targeting HDAC inhibitor bocodepsin/OKI-179 to promote apoptosis. Breast Cancer Research. 2024/03/01 2024;26(1):35. doi:10.1186/s13058-024-01799-5

17. Gao W, Sun L, Gai J, Cao Y, Zhang S. Exploring the resistance mechanism of triple-negative breast cancer to paclitaxel through the scRNA-seq analysis. PloS one. 2024;19(1):e0297260. doi:10.1371/journal.pone.0297260

18. Zhu H, Sarkar S, Scott L, et al. Doxorubicin Redox Biology: Redox Cycling, Topoisomerase Inhibition, and Oxidative Stress. Reactive oxygen species (Apex, NC). 2016;1(3):189–198. doi:10.20455/ros.2016.835

19. Verhaak RGW, Bafna V, Mischel PS. Extrachromosomal oncogene amplification in tumour pathogenesis and evolution. Nat Rev Cancer. May 2019;19(5):283–288. doi:10.1038/s41568-019-0128-6

20. Gu Y, Song Y, Liu J. Identification and characterization of eccDNA-driven genes in humans. PloS one. 2025;20(6):e0324438. doi:10.1371/journal.pone.0324438

21. Ujiie H, Sakyo T, Oya K, et al. Machine Learning-Driven Integration of Cancer Cell Phenotypes Predicts Cisplatin Sensitivity. Cancer medicine. Nov 2025;14(22):e71373. doi:10.1002/cam4.71373

22. Breiman L. Random Forests. Machine Learning. 2001/10/01 2001;45(1):5–32. doi:10.1023/A:1010933404324

23. Chen T, Guestrin C. XGBoost: A Scalable Tree Boosting System. presented at: Proceedings of the 22nd ACM SIGKDD International Conference on Knowledge Discovery and Data Mining; 2016; San Francisco, California, USA. 10.1145/2939672.2939785

24. Zhang Z. Model building strategy for logistic regression: purposeful selection. Annals of translational medicine. Mar 2016;4(6):111. doi:10.21037/atm.2016.02.15

25. Chao C-M, Yu Y-W, Cheng B-W, Kuo Y-L. Construction the Model on the Breast Cancer Survival Analysis Use Support Vector Machine, Logistic Regression and Decision Tree. Journal of Medical Systems. 2014/08/14 2014;38(10):106. doi:10.1007/s10916-014-0106-1

26. Kaur P, Nagaraja GM, Zheng H, et al. A mouse model for triple-negative breast cancer tumor-initiating cells (TNBC-TICs) exhibits similar aggressive phenotype to the human disease. BMC Cancer. Mar 27 2012;12:120. doi:10.1186/1471-2407-12-120

27. Wingett S, Andrews S. FastQ Screen: A tool for multi-genome mapping and quality control [version 2; peer review: 4 approved]. F1000Research. 2018;7(1338)doi:10.12688/f1000research.15931.2

28. Bolger AM, Lohse M, Usadel B. Trimmomatic: a flexible trimmer for Illumina sequence data. Bioinformatics (Oxford, England). Aug 1 2014;30(15):2114–20. doi:10.1093/bioinformatics/btu170

29. Kim D, Paggi JM, Park C, Bennett C, Salzberg SL. Graph-based genome alignment and genotyping with HISAT2 and HISAT-genotype. Nature Biotechnology. 2019/08/01 2019;37(8):907–915. doi:10.1038/s41587-019-0201-4

30. Grabherr MG, Haas BJ, Yassour M, et al. Full-length transcriptome assembly from RNA-Seq data without a reference genome. Nature Biotechnology. 2011/07/01 2011;29(7):644–652. doi:10.1038/nbt.1883

31. Liao Y, Smyth GK, Shi W. featureCounts: an efficient general purpose program for assigning sequence reads to genomic features. Bioinformatics (Oxford, England). 2014;30(7):923–930. doi:10.1093/bioinformatics/btt656

32. Love MI, Huber W, Anders S. Moderated estimation of fold change and dispersion for RNA-seq data with DESeq2. Genome Biology. 2014/12/05 2014;15(12):550. doi:10.1186/s13059-014-0550-8

33. Marini F, Binder H. pcaExplorer: an R/Bioconductor package for interacting with RNA-seq principal components. BMC Bioinformatics. 2019/06/13 2019;20(1):331. doi:10.1186/s12859-019-2879-1

34. Parvizpour S, Razmara J, Pourseif MM, Omidi Y. In silico design of a triple-negative breast cancer vaccine by targeting cancer testis antigens. BioImpacts: BI. 2018;9(1):45.

35. Lawal HA, Uzairu A, Uba S. QSAR, molecular docking studies, ligand-based design and pharmacokinetic analysis on Maternal Embryonic Leucine Zipper Kinase (MELK) inhibitors as potential anti-triple-negative breast cancer (MDA-MB-231 cell line) drug compounds. Bulletin of the National Research Centre. 2021;45:1–20.

36. Prabhavathi H, Dasegowda K, Renukananda K, Karunakar P, Lingaraju K, Raja Naika H. Molecular docking and dynamic simulation to identify potential phytocompound inhibitors for EGFR and HER2 as anti-breast cancer agents. Journal of Biomolecular Structure and Dynamics. 2022;40(10):4713–4724.

37. Akash S, Kumer A, Chandro A, Chakma U, Matin MM. Quantum calculation, docking, ADMET and molecular dynamics of ketal and non-ketal forms of D-glucofuranose against bacteria, black & white fungus, and triple-negative breast cancer. Bioint Res Appl Chem. 2023;13(4):374.

38. Akash S, Aovi FI, Azad MA, et al. A drug design strategy based on molecular docking and molecular dynamics simulations applied to development of inhibitor against triple-negative breast cancer by Scutellarein derivatives. PloS one. 2023;18(10):e0283271.

39. Manivannan HP, Veeraraghavan VP, Francis AP. Identification of molecular targets of Trigonelline for treating breast cancer through network pharmacology and bioinformatics-based prediction. Molecular Diversity. 2023:1–23.

40. Taghizadeh E, Heydarheydari S, Saberi A, JafarpoorNesheli S, Rezaeijo SM. Breast cancer prediction with transcriptome profiling using feature selection and machine learning methods. BMC Bioinformatics. 2022/10/01 2022;23(1):410. doi:10.1186/s12859-022-04965-8

41. Kurian B, Jyothi VL. Breast cancer prediction using ensemble voting classifiers in next-generation sequences. Soft Computing. 2023/06/21 2023;doi:10.1007/s00500-023-08658-z

42. Partin A, Brettin T, Evrard YA, et al. Learning curves for drug response prediction in cancer cell lines. BMC Bioinformatics. 2021/05/17 2021;22(1):252. doi:10.1186/s12859-021-04163-y

43. Sathyanarayanan S, Tantri BR. Confusion Matrix-Based Performance Evaluation Metrics. African Journal of Biomedical Research. 11/30 2024;27(4S):4023-4031. doi:10.53555/AJBR.v27i4S.4345

44. Hazra A. Using the confidence interval confidently. Journal of Thoracic Disease. 2017;9(10):4124–4129.

45. Fawcett T. An introduction to ROC analysis. Pattern Recognition Letters. 2006/06/01/ 2006;27(8):861–874. doi:10.1016/j.patrec.2005.10.010

46. Ribeiro MT, Singh S, Guestrin C. “Why Should I Trust You?”: Explaining the Predictions of Any Classifier. presented at: Proceedings of the 22nd ACM SIGKDD International Conference on Knowledge Discovery and Data Mining; 2016; San Francisco, California, USA. 10.1145/2939672.2939778

47. Murugan TK, Karthikeyan P, Sekar P. Efficient breast cancer detection using neural networks and explainable artificial intelligence. Neural Computing and Applications. 2025/02/01 2025;37(5):3759–3776. doi:10.1007/s00521-024-10790-2

48. Alabi RO, Elmusrati M, Leivo I, Almangush A, Mäkitie AA. Machine learning explainability in nasopharyngeal cancer survival using LIME and SHAP. Scientific Reports. 2023/06/02 2023;13(1):8984. doi:10.1038/s41598-023-35795-0

49. Barredo Arrieta A, Díaz-Rodríguez N, Del Ser J, et al. Explainable Artificial Intelligence (XAI): Concepts, taxonomies, opportunities and challenges toward responsible AI. Information Fusion. 2020/06/01/ 2020;58:82–115. doi:10.1016/j.inffus.2019.12.012

50. Zhu Y, Gujar AD, Wong CH, et al. Oncogenic extrachromosomal DNA functions as mobile enhancers to globally amplify chromosomal transcription. Cancer cell. May 10 2021;39(5):694-707.e7. doi:10.1016/j.ccell.2021.03.006

51. Kim G-E, Kim NI, Lee JS, Park MH, Kang K. Differentially Expressed Genes in Matched Normal, Cancer, and Lymph Node Metastases Predict Clinical Outcomes in Patients With Breast Cancer. Applied Immunohistochemistry & Molecular Morphology. 2020;28(2):111–122. doi:10.1097/pai.0000000000000717

52. Schrors B, Boegel S, Albrecht C, et al. Multi-Omics Characterization of the 4T1 Murine Mammary Gland Tumor Model. Front Oncol. 2020;10:1195. doi:10.3389/fonc.2020.01195

53. Lynch M. Evolution of the mutation rate. Trends in genetics : TIG. Aug 2010;26(8):345–52. doi:10.1016/j.tig.2010.05.003

54. Turner KM, Deshpande V, Beyter D, et al. Extrachromosomal oncogene amplification drives tumour evolution and genetic heterogeneity. Nature. Mar 2 2017;543(7643):122–125. doi:10.1038/nature21356

55. Zeng T, Huang W, Cui L, et al. The landscape of extrachromosomal circular DNA (eccDNA) in the normal hematopoiesis and leukemia evolution. Cell Death Discov. Sep 28 2022;8(1):400. doi:10.1038/s41420-022-01189-w

56. Yin W, Xiang D, Wang T, et al. The inhibition of ABCB1/MDR1 or ABCG2/BCRP enables doxorubicin to eliminate liver cancer stem cells. Sci Rep. May 24 2021;11(1):10791. doi:10.1038/s41598-021-89931-9

57. Karami Fath M, Karimfar N, Fazlollahpour Naghibi A, et al. Revisiting characteristics of oncogenic extrachromosomal DNA as mobile enhancers on neuroblastoma and glioma cancers. Cancer cell international. May 25 2022;22(1):200. doi:10.1186/s12935-022-02617-8

58. Chen Y, Qiu Q, She J, Yu J. Extrachromosomal circular DNA in colorectal cancer: biogenesis, function and potential as therapeutic target. Oncogene. 2023/03/01 2023;42(13):941–951. doi:10.1038/s41388-023-02640-7

59. Hung KL, Jones MG, Wong IT-L, et al. Coordinated inheritance of extrachromosomal DNAs in cancer cells. Nature. 2024/11/01 2024;635(8037):201–209. doi:10.1038/s41586-024-07861-8

60. Lange JT, Rose JC, Chen CY, et al. The evolutionary dynamics of extrachromosomal DNA in human cancers. Nature Genetics. 2022/10/01 2022;54(10):1527–1533. doi:10.1038/s41588-022-01177-x

61. Bergonzini C, Gregori A, Hagens TMS, et al. ABCB1 overexpression through locus amplification represents an actionable target to combat paclitaxel resistance in pancreatic cancer cells. Journal of experimental & clinical cancer research : CR. Jan 2 2024;43(1):4. doi:10.1186/s13046-023-02879-8

62. Hung KL, Yost KE, Xie L, et al. ecDNA hubs drive cooperative intermolecular oncogene expression. Nature. 2021/12/01 2021;600(7890):731–736. doi:10.1038/s41586-021-04116-8

63. Dong Y, He Q, Chen X, Yang F, He L, Zheng Y. Extrachromosomal DNA (ecDNA) in cancer: mechanisms, functions, and clinical implications. Front Oncol. 2023;13:1194405. doi:10.3389/fonc.2023.1194405

64. Abd El-Aziz YS, Spillane AJ, Jansson PJ, Sahni S. Role of ABCB1 in mediating chemoresistance of triple-negative breast cancers. Biosci Rep. Feb 26 2021;41(2)doi:10.1042/bsr20204092

65. Skinner KT, Palkar AM, Hong AL. Genetics of ABCB1 in Cancer. Cancers (Basel). Aug 24 2023;15(17)doi:10.3390/cancers15174236

66. Dong J, Yuan L, Hu C, Cheng X, Qin J-J. Strategies to overcome cancer multidrug resistance (MDR) through targeting P-glycoprotein (ABCB1): An updated review. Pharmacology & Therapeutics. 2023/09/01/ 2023;249:108488. doi:10.1016/j.pharmthera.2023.108488

67. Pilotto Heming C, Muriithi W, Wanjiku Macharia L, Niemeyer Filho P, Moura-Neto V, Aran V. P-glycoprotein and cancer: what do we currently know? Heliyon. Oct 2022;8(10):e11171. doi:10.1016/j.heliyon.2022.e11171

68. Giacomini KM, Huang S-M, Tweedie DJ, et al. Membrane transporters in drug development. Nature Reviews Drug Discovery. 2010/03/01 2010;9(3):215–236. doi:10.1038/nrd3028

69. Pirie R, Stanway-Gordon HA, Stewart HL, et al. An analysis of the physicochemical properties of oral drugs from 2000 to 2022. 10.1039/D4MD00160E. RSC Medicinal Chemistry. 2024;15(9):3125–3132. doi:10.1039/D4MD00160E

70. Yang W, Lipert M, Nofsinger R. Current screening, design, and delivery approaches to address low permeability of chemically synthesized modalities in drug discovery and early clinical development. Drug Discovery Today. 2023/09/01/ 2023;28(9):103685. doi:10.1016/j.drudis.2023.103685

71. Horenko I. On a scalable entropic breaching of the overfitting barrier for small data problems in machine learning. Neural Computation. 2020;32(8):1563–1579.

72. Uddin KMM, Bhuiyan MTA, Saad MN, Islam A, Islam MM. Ensemble Machine Learning-Based Approach to Predict Cervical Cancer with Hyperparameter Tuning and Model Explainability. Biomedical Materials & Devices. 2025/09/01 2025;3(2):1463–1490. doi:10.1007/s44174-024-00268-z

73. Johannssen A, Chukhrova N. The crucial role of explainable artificial intelligence (XAI) in improving health care management. Health care management science. Sep 2025;28(3):565–570. doi:10.1007/s10729-025-09720-y

74. Ponce-Bobadilla AV, Schmitt V, Maier CS, Mensing S, Stodtmann S. Practical guide to SHAP analysis: Explaining supervised machine learning model predictions in drug development. Clinical and translational science. Nov 2024;17(11):e70056. doi:10.1111/cts.70056

75. Zhao S, Macakova K, Sinson JC, et al. Clinical validation of RNA sequencing for Mendelian disorder diagnostics. The American Journal of Human Genetics. 2025/04/03/ 2025;112(4):779–792. doi:10.1016/j.ajhg.2025.02.006

76. Kaya C, Dorsaint P, Mercurio S, et al. Limitations of Detecting Genetic Variants from the RNA Sequencing Data in Tissue and Fine-Needle Aspiration Samples. Thyroid : official journal of the American Thyroid Association. Apr 2021;31(4):589–595. doi:10.1089/thy.2020.0307

77. Zhang A, Miao K, Sun H, Deng CX. Tumor heterogeneity reshapes the tumor microenvironment to influence drug resistance. International journal of biological sciences. 2022;18(7):3019–3033. doi:10.7150/ijbs.72534

78. Fu YC, Liang SB, Luo M, Wang XP. Intratumoral heterogeneity and drug resistance in cancer. Cancer Cell Int. Mar 18 2025;25(1):103. doi:10.1186/s12935-025-03734-w

79. Wu Q, Siddharth S, Sharma D. Triple Negative Breast Cancer: A Mountain Yet to Be Scaled Despite the Triumphs. Cancers (Basel). Jul 23 2021;13(15)doi:10.3390/cancers13153697

80. Luo J, Li Y, Zhang T, et al. Extrachromosomal circular DNA in cancer drug resistance and its potential clinical implications. Front Oncol. 2022;12:1092705. doi:10.3389/fonc.2022.1092705

81. Kim H, Kim S, Wade T, et al. Mapping extrachromosomal DNA amplifications during cancer progression. Nature Genetics. 2024/11/01 2024;56(11):2447–2454. doi:10.1038/s41588-024-01949-7

82. Vyshnavi H, Namboori K. Identifying potential ligand molecules EGFR mediated TNBC targeting the kinase domain-identification of customized drugs through in silico methods. Research in pharmaceutical sciences. Apr 2023;18(2):121–137. doi:10.4103/1735-5362.367792

83. Steiner T, R. Desiraju G. Distinction between the weak hydrogen bond and the van der Waals interaction. 10.1039/A708099I. Chemical Communications. 1998;(8):891–892. doi:10.1039/A708099I

84. Chen D, Oezguen N, Urvil P, Ferguson C, Dann SM, Savidge TC. Regulation of protein-ligand binding affinity by hydrogen bond pairing. Science advances. Mar 2016;2(3):e1501240. doi:10.1126/sciadv.1501240

85. Gottesman MM, Fojo T, Bates SE. Multidrug resistance in cancer: role of ATP-dependent transporters. Nat Rev Cancer. Jan 2002;2(1):48–58. doi:10.1038/nrc706

86. Aller SG, Yu J, Ward A, et al. Structure of P-glycoprotein reveals a molecular basis for poly-specific drug binding. Science. Mar 27 2009;323(5922):1718–22. doi:10.1126/science.1168750

87. Jaworski JJ, Pfuderer PL, Czyz P, Petris G, Boemo MA, Sale JE. ecDNA replication is disorganized and vulnerable to replication stress. Nucleic acids research. Jul 19 2025;53(14)doi:10.1093/nar/gkaf711

